# Control of Hematopoietic Stem and Progenitor Cell Function Through Epigenetic Regulation of Energy Metabolism and Genome Integrity

**DOI:** 10.1101/554519

**Authors:** Zhenhua Yang, Kushani Shah, Alireza Khodadadi-Jamayran, Hao Jiang

## Abstract

It remains largely unclear how stem cells regulate bioenergetics and genome integrity to ensure tissue homeostasis. Here, our integrative gene analyses suggest metabolic and genotoxic stresses may underlie the common functional defects of both fetal and adult hematopoietic stem and progenitor cells (HSPCs) upon loss of DPY30, an epigenetic modulator that facilitates H3K4 methylation. DPY30 directly regulates expression of several key glycolytic genes, and its loss in HSPCs critically impaired energy metabolism, including both glycolytic and mitochondrial pathways. We also found significant increase in DNA breaks as a result of impaired DNA repair upon DPY30 loss, and inhibition of DNA damage response partially rescued clonogenicity of the DPY30-deficient HSPCs. Moreover, CDK inhibitor *p21* was upregulated in DPY30-deficient HSPCs, and *p21* deletion alleviated their functional defect. These results demonstrate that epigenetic mechanisms by H3K4 methylation play a crucial role in HSPC function through control of energy metabolism and protecting genome integrity.

## INTRODUCTION

Adult tissue stem cells, including hematopoietic stem and progenitor cells (HSCs and HPCs, or HSPCs), usually remain in the quiescent state with limited cell divisions, but can enter rapid proliferation upon activation (Kondo et al., 2003). As one of the most fundamental processes of all lives, metabolism profoundly influences the fate determination of stem and progenitor cells (Ito and Suda, 2014). Quiescent HSCs have relatively inactive mitochondria and mainly rely on glycolysis, but rapidly switch to mitochondrial oxidative phosphorylation as the major energy supply when they undergo activation and differentiation (Ito and Suda, 2014). Regulatory mechanisms in gene expression, including epigenetic mechanisms, can play an important role in stem and progenitor cell fate determination through impinging on the expression of rate-limiting enzymes in the metabolic pathways. Histone H3K4 methylation is one of the most prominent epigenetic marks associated with active gene expression (Shilatifard, 2008), but its role in regulation of metabolism, especially in stem and progenitor cell settings, remains unclear.

Maintaining the stability of the genetic material through the intricate network of DNA damage response (DDR) is particularly crucial for stem and progenitor cells to ensure the propagation of correct genetic information throughout the daughter cells in the tissue for the benefit of the whole organism (Blanpain et al., 2011). Mice deficient of components in either recognition (Ito et al., 2004) or repair (Nijnik et al., 2007; Rossi et al., 2007) of DNA lesions often have defective HSCs. As DDR occurs in the context of chromatin, it is thus governed by epigenetic mechanisms regulating chromatin structure and accessibility (Lukas et al., 2011). In addition to the well-established association with active gene expression, a role of H3K4 methylation in regulating genome stability is also gradually emerging (Burman et al., 2015; Faucher and Wellinger, 2010; Herbette et al., 2017; Higgs et al., 2018).

The most notable H3K4 methylation enzymes in mammals are the SET1/MLL complexes (Shilatifard, 2008), which comprise one of six different catalytic subunits and several shared core subunits including DPY30. DPY30 directly facilitates genome-wide H3K4 methylation (Jiang et al., 2011) and plays important roles in the fate transition between pluripotent and differentiated states (Jiang et al., 2011; Yang et al., 2015). We have further shown a crucial role of DPY30 in HSPCs and animal hematopoiesis (Yang et al., 2014). Ablation of *Dpy30* in the hematopoietic system of the adult mouse bone marrow (BM) leads to loss of global H3K4 methylation, and disables differentiation and long-term maintenance of adult HSCs (Yang et al., 2016). This is shown in part by the striking accumulation of phenotypic HSCs and early HPCs at the expense of more downstream hematopoietic cells after *Dpy30* ablation in BM (Yang et al., 2016), a phenotype shared by loss of other subunits of the SET1/MLL complexes (Arndt et al., 2018; Chen et al., 2014; Chun et al., 2014; Santos et al., 2014). Moreover, DPY30-deficient adult HSCs contribute poorly to all hematopoietic cell populations in the later stage after transplant. The normal transition of gene program between cell populations is also disrupted, and multiple genes important for HSC maintenance and differentiation are dysregulated following DPY30 loss (Yang et al., 2016). However, it is unclear what pathways are fundamentally important for DPY30’s role in HSPC fate determination.

As fetal and adult HSCs differ in many aspects including cell cycle regulation, self-renewal potential, surface marker expression and reconstitution ability (Orkin and Zon, 2008), the function of DPY30 and H3K4 methylation in prenatal HSCs is still unknown. We reason that pathways fundamentally important for the activity of an epigenetic modulator are likely to be conserved in different biological systems. In this work, we first revealed a similar requirement of DPY30 in the activation and long-term maintenance of fetal HSPCs as compared to the adult HSPCs. Our dissection of targets shared in these two systems allowed us to identify energy metabolism and DDR as fundamentally important mediators of the H3K4 methylation pathway in hematopoiesis, as further supported by our rescue assays.

## RESULTS

### DPY30 deficiency in fetal liver results in anemia and accumulation of early HSPCs

We previously generated a *Dpy30* conditional knockout (KO) mouse model where the Floxed (*F*) allele of *Dpy30* is converted to a null allele upon Cre activity (Yang et al., 2016). Based on this and the *Vav-Cre* (Stadtfeld and Graf, 2005) mouse models, here we generated *Dpy30*^*F/+*^, *Dpy30*^*F/-*^, *Vav-Cre; Dpy30*^*F/+*^, and *Vav-Cre; Dpy30*^*F/-*^ fetuses in the same litters. The *Vav-Cre; Dpy30*^*F/-*^ fetuses developed normally until E13.5, a time when Vav-Cre-mediated excision in HSCs becomes fully penetrant (Gan et al., 2010), but were never born and thus likely died at the late embryonic stage. The *Vav-Cre; Dpy30*^*F/-*^ fetuses were anemic at E14.5 and anemia became more severe at E15.5 (**Figure 1A**). No difference was observed among the *Dpy30*^*F/+*^, *Dpy30*^*F/-*^, and *Vav-Cre; Dpy30*^*F/+*^ fetuses. We confirmed *Dpy30* excision (**Figure S1A**) and dramatic reduction of *Dpy30* mRNA (**Figure 1B**) in total and various HSPC populations (see **Table S1** for cell surface markers) of the *Vav-Cre; Dpy30*^*F/-*^ fetal liver (FL), and reduction of the DPY30 protein (**Figure 1C**) in total FL. DPY30 loss also resulted in marked reduction in H3K4 tri-methylation (H3K4me3) and mild reduction in H3K4 mono- and di-methylation (**Figure 1C**).

**Figure 1.**
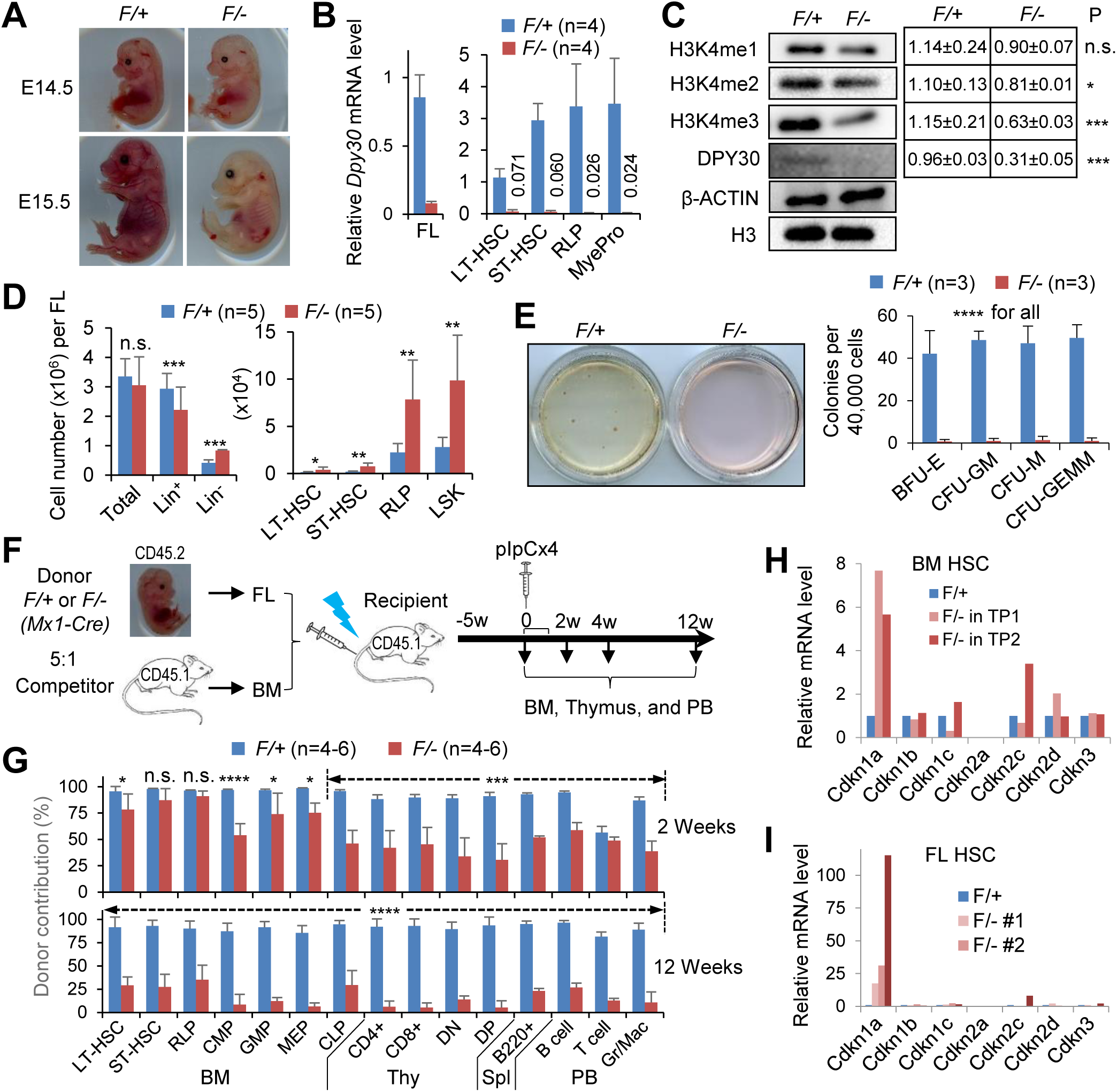
DPY30 deficiency in the fetal hematopoietic system results in anemia and defective HSC function. *Vav-Cre; Dpy30*^*F/+*^ (*F*/+) and *Vav-Cre; Dpy30*^*F/-*^ (*F*/-) littermate fetuses were used in panels (A-E) and (I). **(A)** Image of littermate fetuses at E14.5 and E15.5. **(B)** Relative *Dpy30* mRNA levels in whole FL and sorted hematopoietic cell populations determined by RT-qPCR and normalized to *Actb*. n=3-6 each for FL, n=4 each for sorted cells. **(C)** Immunoblotting for different levels of H3K4 methylation and other proteins in FL. Right, quantification from 5 embryos each. **(D)** Absolute numbers of FL cell populations. n=5 each. **(E)** Colony formation assay using FL cells, showing representative images of colonies and quantification. n=3 each. **(F)** Scheme for the mixed chimera transplantation system using whole FL cells from *Mx1-Cre; Dpy30*^*F/+*^ or *Mx1-Cre; Dpy30*^*F/-*^ fetuses as donors and whole BM cells from wild type mice as competitors. **(G)** Donor contribution to different cell populations in chimeras at indicated times after pIpC injections following scheme in (G). n=4-6 each. **(H and I)** Relative expression of CDK inhibitor genes in control (*F/+*) versus *Dpy30* KO (*F/-*) HSCs in two independent BM chimeras (H) and three FLs at E14.5 (I), as analyzed by RNA-seq. The expression levels 17 in *F/+* cells were set as 1. The BM HSC data are based on RNA-seq results in donor (*Mx1-Cre; Dpy30*^*F/+*^ or *Mx1-Cre; Dpy30*^*F/-*^)-derived HSCs in the BM chimera recipients in two independent BM transplantations (TP1 and TP2) from our previous work (Yang et al., 2016). Data are shown as mean ±SD for (C-E). *P<0.05, **P<0.01, ***P<0.001, ****P<0.0001, by 2-tailed Student’s *t*-test. See also Figures S1-S2.

While the number of total mononuclear FL cells was unaffected, we consistently detected a striking accumulation (**Figures 1D, S1B, and S1C**), in absolute number and frequency, of lineage negative (Lin^-^) cells and early HSPCs including LT-HSCs, ST-HSCs, Restricted Lineage Progenitors (RLPs) (Kiel et al., 2005), and Lin^-^ Sca1^+^c-Kit^+^ (LSK) cells in the *Vav-Cre; Dpy30*^*F/-*^ FLs. Conversely, the frequency of lineage positive (Lin^+^) cells was decreased (**Figure S1B**). DPY30 loss in FL did not affect proliferation of multipotent stem cells (LT-HSCs and ST-HSCs), but significantly reduced that of oligopotent progenitors (RLP) and Lin^+^ cells (**Figure S1D, left**). Apoptosis was unaffected in all FL cell fractions upon DPY30 loss (**Figure S1D, right**).

### DPY30-deficient fetal liver HSCs are defective in differentiation and long-term maintenance

Compared to *Vav-Cre*; *Dpy30*^*F/+*^, the *Vav-Cre*; *Dpy30*^*F/-*^ FL cells were severely defective in forming any types of colonies (**Figure 1E**), and in reconstituting multiple peripheral blood (PB) lineages in competitive transplantation assays, with much lower contributions to hematopoietic cells at all stages (**Figures S1E-S1G**). We also established a mixed FL-BM chimeras assay and induced *Dpy30* excision by injection of Polyinosinic-polycytidylic acid (pIpC) into recipients after stable engraftment of the donor FLs carrying *Mx1-cre* (Yang et al., 2016) (**Figure 1F**). At two weeks after pIpC injection, *Dpy30* deletion had mild or no effects on donor contribution to early HSPCs, but significantly reduced that to oligopotent HPCs, developing T cells in thymus and B cells in spleen, and mature blood cells in PB (**Figure 1G**), consistent with a multi-lineage differentiation defect. At twelve weeks after pIpC injection, we detected a dramatic reduction in the contribution of the KO donor cells to all cell populations including HSCs (**Figure 1G**). Taken together, these results demonstrated an essential role of DPY30 for FL HSC differentiation and long-term maintenance, similar to its role in adult hematopoiesis (Yang et al., 2016).

### Differential and common regulation of gene expression in fetal liver and adult bone marrow HSCs upon *Dpy30* ablation

The largely similar phenotypes of the *Dpy30* KO FL and BM HSCs suggest that DPY30 and certain DPY30 targets are fundamentally important in regulating HSCs regardless of the developmental stages. However, we found that DPY30 loss in FLs did not reduce the expression of some key regulatory genes for adult HSC differentiation and maintenance (Yang et al., 2016) (**Figure S2A**). As shown by RNA-seq analysis on FL LT-HSCs (**Table S2**), genes downregulated upon DPY30 loss in FLs were enriched in many different functions, and those upregulated were enriched in membrane proteins and immune functions (**Figure S2B**). In the 275 and 577 genes downregulated over two fold upon DPY30 loss in FL and BM HSCs, respectively, only 21 genes were shared, with no significant enrichment of pathways (**Figure S2C and Table S2**). The number of shared genes increased to 95 when we used 1.5-fold cutoff, with an enrichment of nucleotide excision repair genes and mitochondrial genes (**Figure S2E and Table S2**). Meanwhile, in the 340 and 669 genes that were upregulated over two fold upon DPY30 loss in FL and BM HSCs, respectively, 41 genes were shared (**Figure S2D and Table S2**). The number of shared genes increased to 138 when we used 1.5-fold cutoff, with the most significant enrichment in inflammation-related genes (**Figures S2D and S2F**). This suggests cellular stresses upon loss of DPY30 in both FL and BM HSCs, which is also reflected by the specific and dramatic upregulation of *Cdkn1a* (*p21*) in both types of HSCs as well as BM myeloid progenitors (**Figures 1H, 1I, and S2G**), a gene well-known to be induced by a wide spectrum of stress stimuli including genotoxic, metabolic, and oxidative insult (Gorospe et al., 1999). Indeed, several genes involved in redox homeostasis, such as *Alox5, Cybb* (*Nox2*), and *Xdh*, were commonly upregulated in both types of HSCs upon DPY30 loss (**Table S2**). Together with the enrichment of DNA repair and mitochondrial genes being commonly downregulated upon DPY30 loss, these results prompted us to further study the effect of DPY30 loss on the metabolic state and DNA integrity of HSCs.

### DPY30 is important for glucose metabolism in HSPCs through direct regulation of key glycolytic genes

We efficiently deleted *Dpy30* in various populations of hematopoietic cells (including HSCs) in the BM of *Mx1-Cre*; *Dpy30*^*F/-*^ mice (**Figure S3A**). Consistent with metabolic stress, both immunoblotting and flow cytometry analyses showed a strong increase in phosphorylation of AMPK at Thr172, a well-established sensor and responder of low nutrient or energy (Lin and Hardie, 2018), in all different hematopoietic cell populations following DPY30 loss (**Figures 2A and 2B**). Moreover, phosphorylation of ACETYL-COA CARBOXYLASE (ACC) at Ser79, one of AMPK’s substrates upon AMPK activation, also increased in DPY30-deficient BM (**Figure 2A**). These results suggest an activation of the cellular sensor of low nutrient or energy in response to DPY30 deficiency.

**Figure 2.**
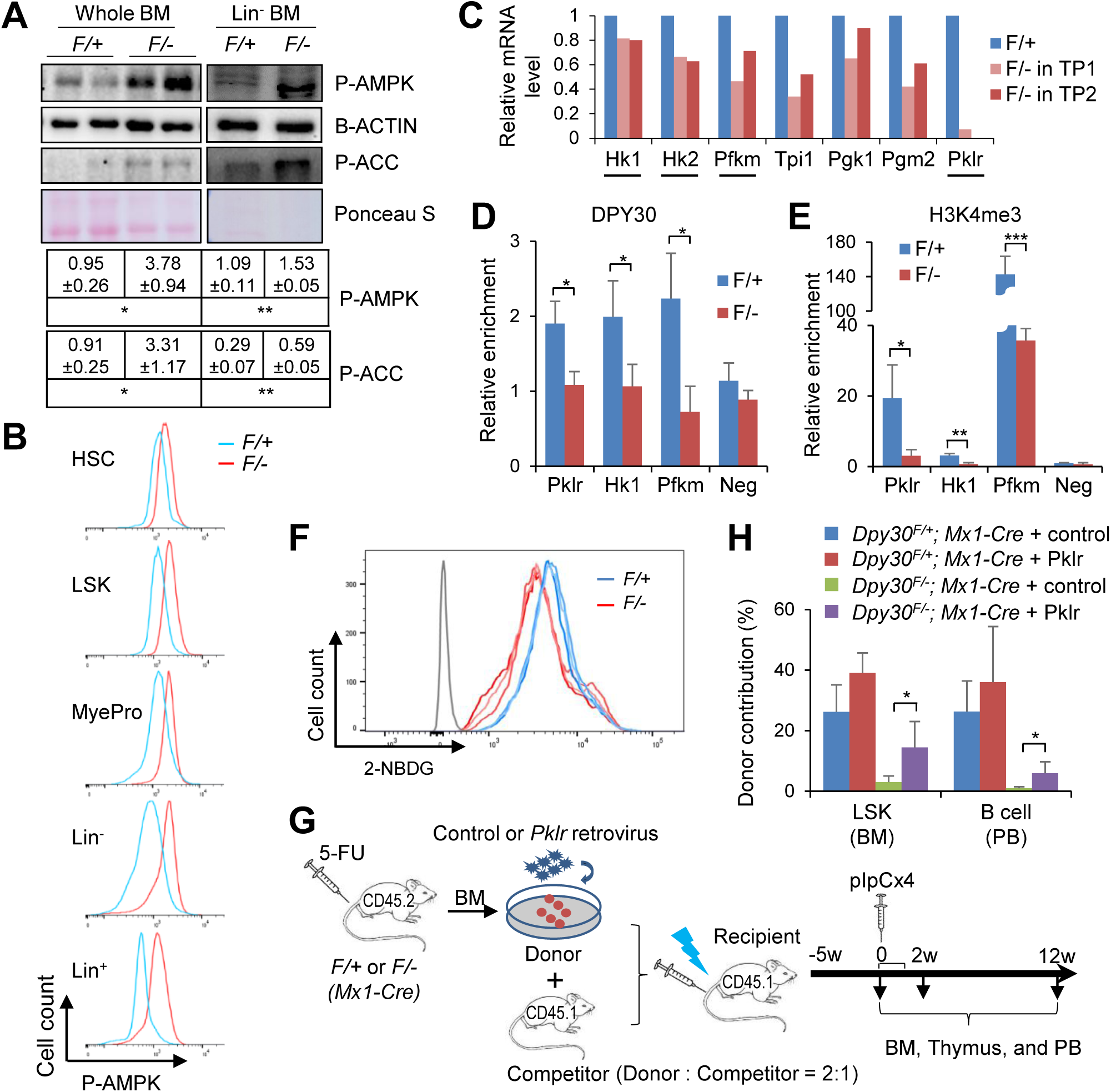
DPY30 deficiency impairs glucose metabolism in BM HSPCs. **(A)** Immunoblotting in whole or Lin^-^ BMs from pIpC-injected *Mx1-Cre; Dpy30*^*F/+*^ and *Mx1-Cre; Dpy30*^*F/-*^ mice. Each lane represents an individual pool of cells. Bottom, quantification from 3 animals each. **(B)** Representative FACS analyses for phospho-AMPK (Thr172) in different cell populations in BMs from pIpC-injected *Mx1-Cre; Dpy30*^*F/+*^ and *Mx1-Cre; Dpy30*^*F/-*^ mice. **(C)** Relative expression of glycolytic genes, as analyzed by RNA-seq in LT-HSCs from two independent BM transplantations from previous work (Yang et al., 2016). Underlined genes are rate-limiting for glycolysis. **(D and E)** ChIP for DPY30 (D) and H3K4me3 (E) using Lin^-^ BM cells from pIpC-injected *Mx1-Cre; Dpy30*^*F/+*^ and *Mx1-Cre; Dpy30*^*F/-*^ mice. n=3 mice each. **(F)** Representative histogram showing reduced 2-NBDG uptake in vitro by sorted LSK cells from pIpC-injected *Mx1-Cre; Dpy30*^*F/+*^ and *Mx1-Cre; Dpy30*^*F/-*^ mice. The gray line indicates isotype control. Each colored line represents an individual mouse out of 3 mice each. **(G)** Scheme for the rescue assay using BMs from *Mx1-Cre; Dpy30*^*F/+*^ (*F/+*) and *Mx1-Cre; Dpy30*^*F/-*^ (*F/-*) mice. Both the control and *Pklr* viral constructs expressed GFP. **(H)** Donor contribution to indicated cell populations in the transplant recipients 2 weeks after pIpC-injections. n=4 mice each. Data are shown as mean ±SD for (A), (D), (E), and (H). *P<0.05, **P<0.01, ***P<0.001, by 2-tailed Student’s *t*-test for (A), (D), and (E), and by 1-factor ANOVA with post hoc *t* test for (H). See also Figures S3.

We then examined the impact of deficiency of DPY30 and H3K4 methylation on glucose metabolism, a major constituent of cellular energy pathway. DPY30 loss in BM and FL HSCs resulted in downregulation of several genes encoding key glycolytic enzymes, including a number of rate-limiting ones, such as HEXOKINASE (HK1 and HK2), PHOSPHOFRUCTOKINASE MUSCLE (PFKM), and PYRUVATE KINASE LIVER AND RBC (PKLR) (**Figures 2C, S3B, and S3C**). DPY30 and H3K4me3 were enriched at the transcription start sites of these genes in the Lin^-^ BM in the control mice, but both signals were significantly reduced in the KO littermates (**Figures 2D and 2E**). These results indicate that DPY30, likely through its activity in facilitating H3K4 methylation, directly regulates the expression of these key glycolytic genes.

HEXOKINASE phosphorylates glucose to form glucose 6-phosphate. This reaction is important in promoting continuous uptake of glucose into the cell, as it keeps the glucose concentration low and also prevents glucose from diffusing out of the cell due to the added charge of the reaction product. We thus predicted that downregulation of *Hk1* and *Hk2* would impair glucose uptake. Indeed, as measured by glucose analog (2-NBDG) uptake by sorted LSK cells, we found that glucose uptake was consistently impaired in HSPCs by DPY30 deficiency (**Figure 2F**). This impairment is probably not a result of reduced glucose transporter expression, as we did not find any significant downregulation of known glucose transporter genes (**Table S2**).

### Transient and modest rescue of hematopoiesis by restoring *Pklr* expression in *Dpy30* KO mice

PKLR catalyzes the transphosphorylation of phosphoenolpyruvate into ATP and pyruvate, a key intermediate in multiple metabolic pathways. Genetic alterations of PKLR are the common cause of chronic hereditary nonspherocytic hemolytic anemia (Zanella et al., 2005). To determine if the profound loss of H3K4me3 and expression of *Pklr* in the *Dpy30* KO HSCs may contribute to the defective hematopoiesis, we attempted to rescue the in vivo HSC activity by overexpressing *Pklr* (**Figures 2G, S3D, and S3E**). *Pklr* overexpression did not affect the donor contribution to any hematopoietic cell populations before *Dpy30* was deleted, but significantly (and partially) rescued the contribution of the *Dpy30*-deleted donors to early HSPCs (BM LSK) and B cells in PB 2 weeks after *Dpy30* deletion (**Figures 2H and S3F**). However, at 12 weeks after *Dpy30* deletion, we failed to observe any rescue for any cell populations (**Figure S3F**). These results suggest that regulation of *Pklr* expression by DPY30 functionally contributes to, but is far from the entirety of, the hematopoietic control by DPY30.

### DPY30 regulates oxidative energy metabolism of the stem cells

Considering the dysregulation of genes involved in redox homeostasis, we next examined the impact of DPY30 deficiency on oxidative energy metabolism. Real-time measurement of oxygen consumption showed that DPY30 deficiency significantly reduced both the basal and the total oxidative capacity of Lin^-^ progenitors (**Figure 3A**), as well as the extracellular acidification rate (**Figure 3B**). We also found that DPY30 deficiency in early HSPCs, including LT-HSCs, ST-HSCs, and RLPs, led to substantial reduction in mitochondrial membrane potential without affecting the mitochondrial mass or DNA level (**Figures 3C-3F and S4A-4C**). Interestingly, DPY30 loss in the lineage-committed BM cells led to marked increase in mitochondrial membrane potential and mass (**Figures S4A and S4B**). Consistent with these results, the level of reactive oxygen species (ROS) was significantly reduced upon DPY30 loss in early HSPCs including HSCs, but increased in more downstream cells (**Figures 3F and S4C**). Partially consistent with BM cells, FL cells showed significant reduction in ROS levels upon DPY30 loss in all different cell populations including early HSPCs and Lin^+^ cells (**Figure S4D**), while primary mouse embryonic fibroblasts (MEFs) showed minimal alteration in ROS level upon DPY30 loss (**Figure S4E**). All these results indicate that DPY30 plays an important role in maintaining the activity of mitochondria and the oxidative energy metabolism. We found that the cellular ATP level was not significantly affected by DPY30 deficiency (**Figure 3G**). While NADP/NADPH ratio was unaffected, NAD^+^/NADH ratio was modestly but significantly increased (**Figure 3H**), suggesting a dysregulated redox homeostasis in DPY30-deficient HSPCs.

**Figure 3.**
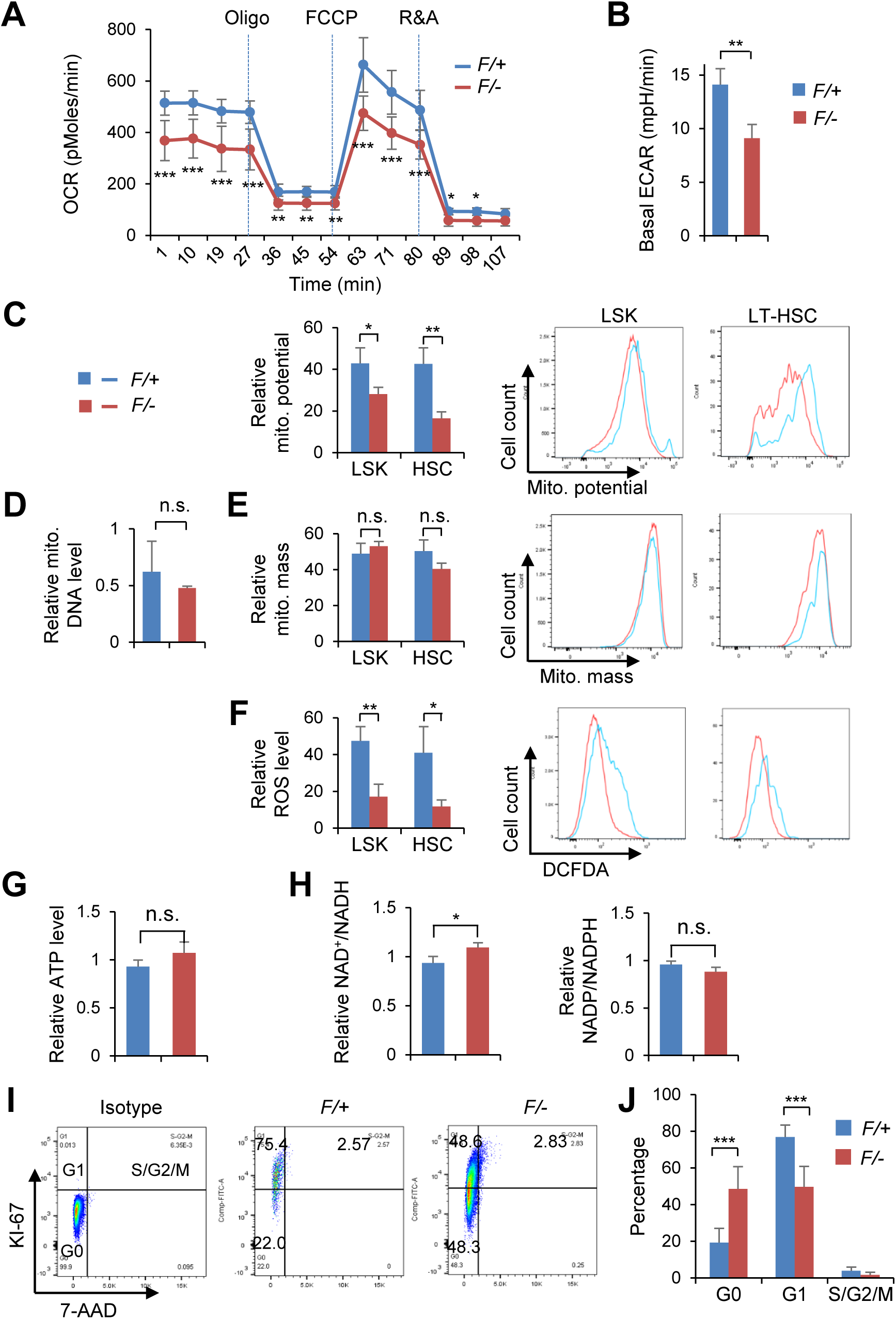
DPY30 deficiency impairs mitochondrial function and alters quiescence state of the BM HSCs. BMs from pIpC-injected *Mx1-Cre; Dpy30*^*F/+*^ (*F/+*) and *Mx1-Cre; Dpy30*^*F/-*^ (*F/-*) mice were used in this figure. **(A and B)** Oxygen consumption rate (OCR) (A) and basal extracellular acidification rates (B) of live Lin^-^ BM cells. n=3 mice each. **(C, E, and F)** Analyses (left) and representative histograms (right) for mitochondrial membrane potential (C), mitochondrial mass (E) and ROS level (F) of indicated BM cell populations. n=3 mice each. **(D)** Mitochondrial DNA levels from BM LSK cells were determined by qPCR on *mt-Co2* and normalized to *Hbb* for genomic DNA levels. n=3 mice each. **(G and H)** Total cellular ATP levels (G), and NAD^+^/NADH and NADP^+^/NADPH ratios (H) in BM LSK cells were determined. n=3 mice each. **(I)** Representative FACS plots of KI-67 staining on LT-HSCs. **(J)** G0, G1, and S/G2/M phases of LT-HSCs were quantified. n = 5 mice each. Data are shown as mean ±SD for all panels. n.s. not significant, *P<0.05, **P<0.01, ***P<0.001, by 2-tailed Student’s *t*-test. See also Figure S4.

To examine the functional relevance of ROS increase in the lineage-committed BM cells to pancytopenia of the DPY30-deficient animals, we administered N-acetyl-cysteine (NAC), a potent antioxidant, into these animals. We found that while NAC only slightly but insignificantly improved the survival of KO mice (**Figure S4F**), it significantly (yet partially) and selectively rescued the concentration of the red blood cells in PB of the KO mice (**Figure S4G**). We also induced ex vivo *Dpy30* deletion in isolated Lin^+^ BM, and found that NAC treatment in cultured BM ameliorated the apoptotic response of erythroid cells that was increased by *Dpy30* deletion, but did not affect the proliferation of Lin^+^ cells that was reduced by Dpy30 depletion (**Figure S4H**). These results suggest that increase of ROS level in the lineage-committed cells functionally contributes to the reduced erythroid cell production upon DPY30 loss.

### DPY30 regulates the quiescent state of the stem cells

Considering the intimate connection of HSC energy state and its cell cycle regulation, we further studied how DPY30 deficiency might affect the HSC cell cycle. We previously showed that DPY30 deficiency did not significantly affect the proliferative capacity of HSCs or LSK cells as measured by short-time incorporation of nucleotide analog BrdU (Yang et al., 2016). Here we analyzed the proliferative state of the HSCs by staining of KI-67, a nuclear protein not only strictly associated with cell proliferation (Scholzen and Gerdes, 2000) but also functionally crucial for successful cell division (Cuylen et al., 2016). DPY30 deficiency resulted in significant increase in the percentage of HSCs in G0 stage but not in the S/G2/M of the cell cycle (**Figures 3I and 3J**), suggesting that DPY30-deficient HSCs were in an aberrant and more deeply quiescent state of cell cycle compared to the control HSCs.

### DPY30 and H3K4 methylation are important for genome integrity of cells

We then focused on genotoxic stress upon DPY30 loss. We found a significant increase in the γ-H2AX level in the FL and Lin^-^ BM cells upon DPY30 loss in vivo (**Figure 4A**), suggesting breach of genome integrity and increase in DDR. We also directly demonstrated a significant increase in DNA breaks following DPY30 loss in early BM HSPCs by using the comet assay (**Figures S5A and S5B**). Since the major source of DNA damage, ROS, was not increased in *Dpy30* KO HSCs (**Figure 3F**), we sought to determine if the increase in DNA damage could be a result of poor resolution of damage in the absence of DPY30. To avoid complication by non-cell-autonomous and cumulative effects for cells derived from the KO mice, we first used primary MEFs from *Dpy30*^*F/F*^ and *CAG-CreER; Dpy30*^*F/F*^ mice. Treatment of the latter cells with 4-OH tamoxifen induced *Dpy30* excision and led to great reduction in *Dpy30* mRNA level (**Figure S5C**) and global H3K4 methylation (**Figure S5E** and immunoblotting results shown before (Yang et al., 2018)). A high level of nuclear γ-H2AX foci was observed in all cells at 15 minutes after ionizing radiation and then gradually reduced to the background level by 24 hours, reflecting the process of DNA repair after damage (**Figures S5D and S5E**). We found that the *Dpy30*-deleted MEFs accumulated similar level of DNA damage as the control MEFs at 15 minutes after radiation, but exhibited significantly higher levels of DNA damage than the control at 2 and 24 hours after radiation (**Figures S5D and S5E**). We then performed similar assays on HSPCs (**Figures 4B-4G, S5F, and S5G**), and monitored DDR and DNA damage levels using nuclear γ-H2AX foci and comet assays (**Figure 4B**). Alternatively, we also irradiated the *Dpy30*-deleted Lin^-^ BM cells and monitored ATM activation reflected by the level of the phosphorylated KAP1 (an ATM substrate) (**Figure 4F**). Compared to control HSPCs, the *Dpy30*-deleted HSPCs had significantly higher levels of DNA breaks (by comet assays, **Figure 4E and S5G**) as well as DDR (by nuclear γ-H2AX foci, **Figures 4D and S5F**, and phosphorylated KAP1, **Figure 4G**) both before and after irradiation. These results indicate that DNA repair capacity is significantly impaired in cells deficient of DPY30 and H3K4 methylation, thus leading to sustained DNA damage and DDR.

**Figure 4.**
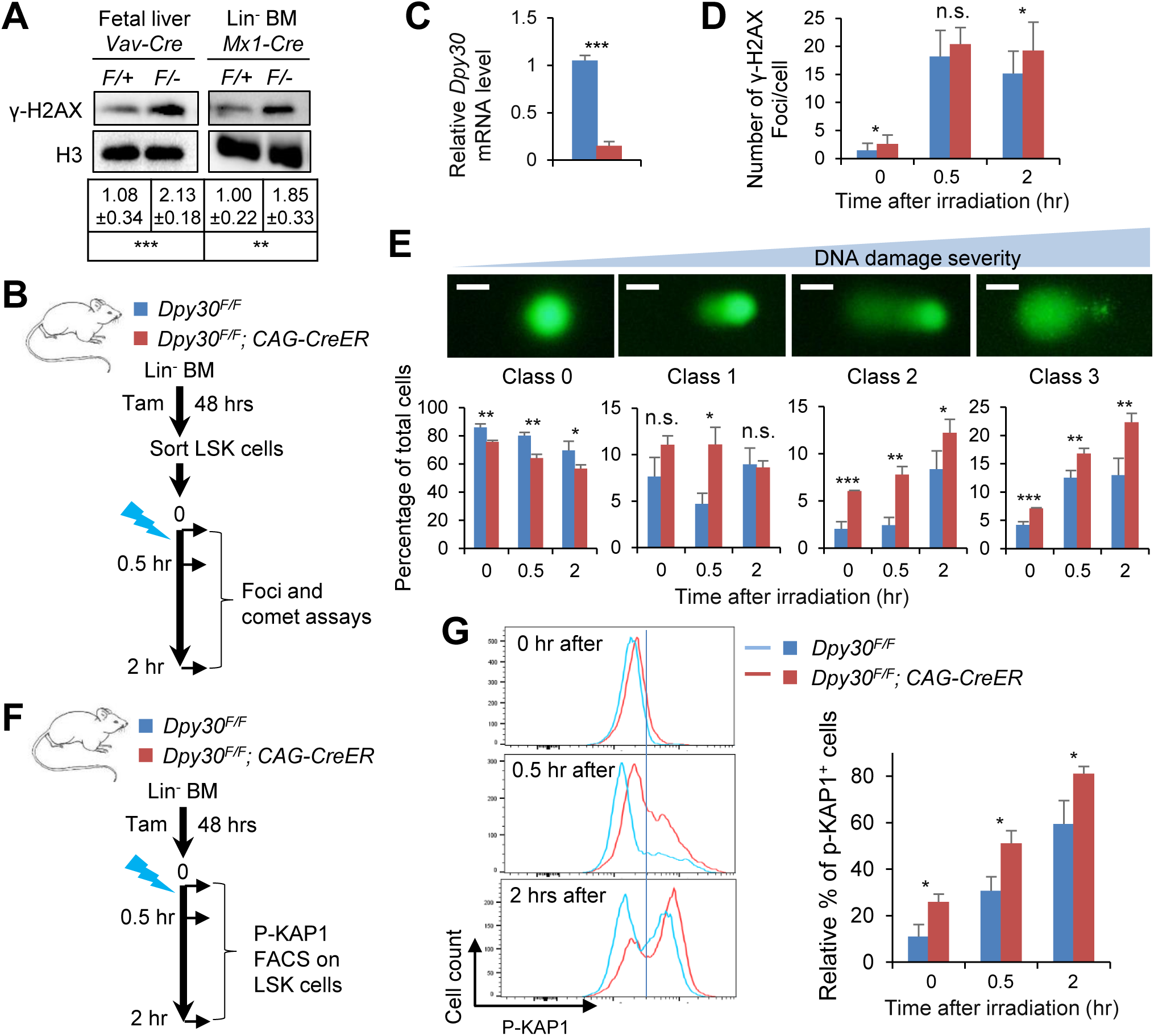
DPY30 deficiency results in increase in DNA damage and impairs DNA repair. **(A)** Immunoblotting of γ-H2AX and H3 in *Vav-Cre; Dpy30*^*F/+*^ and *Vav-Cre; Dpy30*^*F/-*^ FL and Lin^-^ BM cells from pIpC-injected *Mx1-Cre; Dpy30*^*F/+*^ and *Mx1-Cre; Dpy30*^*F/-*^ mice. Bottom, quantification from 5 animals each. **(B)** Scheme for DNA damage assays and the mouse genotype legend for (C-E). LSK cells sorted from tamoxifen-treated Lin^-^ BM were irradiated for assays at different time points. n=3 mice each. **(C)** Relative *Dpy30* mRNA levels were determined by RT-qPCR and normalized to *Actb.* **(D)** γ-H2AX foci numbers per cell from 20-30 LSK cells each. Representative images are in Figure S5F. **(E)** Comet assay on LSK cells. We used 4 classes of cell morphology to show increasing DNA damage severity, and quantified (bottom) the percentages of each class before and after irradiation. **(F)** Assay scheme and the mouse genotype legend for (G). Tamoxifen-treated Lin^-^ BM were irradiated, followed by FACS assays gated on LSK cells at different time points. n=3 mice each. **(G)** FACS analyses for phosphorylated Kap-1 in LSK cells. Left, representative plots. Right, quantification based on the demarcation of the vertical line shown in the left plots. Data are shown as mean ±SD. *P<0.05, **P<0.01, ***P<0.001, by 2-tailed Student’s *t*-test. Scale bars, 10 µm. See also Figure S5.

### Inhibition of DDR partially rescues the function of the DPY30-deficienct HSPCs

We first confirmed that 4-OH-tamoxifen treatment of *CAG-CreER; Dpy30*^*F/F*^ Lin^-^ BM cells greatly reduced their *Dpy30* expression (**Figure 5A**) and capacity in forming all different types of colonies (**Figures 5B and 5C**). We then showed that, while the ATM inhibitor (KU55933) treatment had no effect on the clonogenicity of the control cells, it partially but significantly rescued the ability of the *Dpy30*-deleted cells in forming all different types of colonies (**Figures 5C-5E**). Moreover, pharmacologic inhibition of ATR and CHK1, but not CHK2, also significantly enhanced the clonogenicity of the *Dpy30*-deleted Lin^-^ BM cells in a dose-dependent manner (**Figure S6**). These results suggest that the sustained DDR functionally affects HSPC activities.

**Figure 5.**
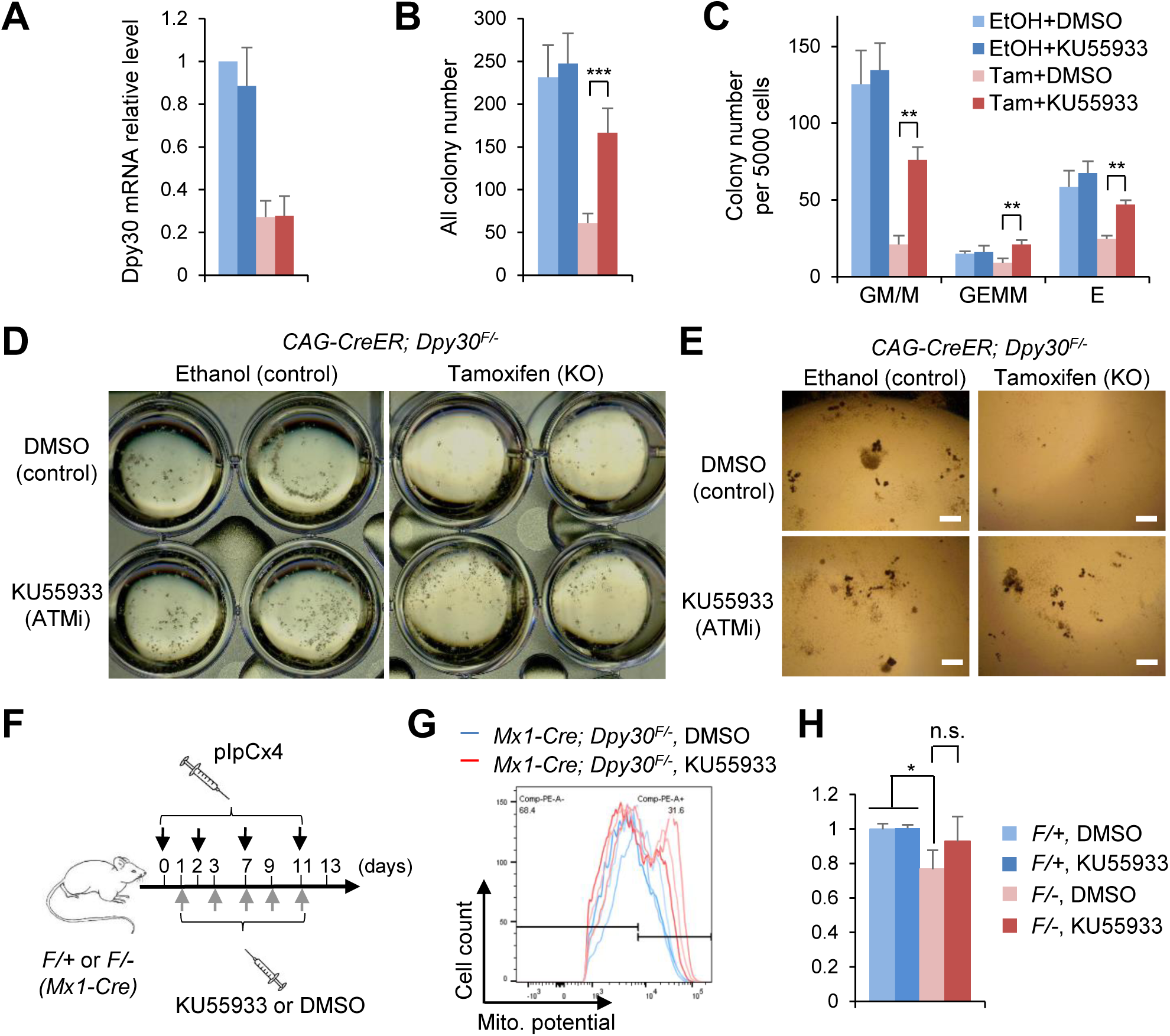
Inhibition of DDR partially rescues the function of the DPY30-deficient hematopoietic cells. **(A-E)** Lin^-^ BM cells from *CAG-CreER; Dpy30*^*F/F*^ mice were treated with indicated agents in liquid culture for two days and subjected to colony formation assays. **(A)** Relative *Dpy30* mRNA levels were determined by RT-qPCR and normalized to *Actb.* n=3 each. **(B and C)** All colonies (B) (3 independent biological repeats) and different types of colonies (C) (2 independent biological repeats) were quantified. **(D and E)** Representative image of the plates (D) and colonies (E) are shown. **(F-H)** *Mx1-Cre; Dpy30*^*F/+*^ *or Mx1-Cre; Dpy30*^*F/-*^ mice were injected with pIpC and either vehicle (DMSO) or ATM inhibitor KU55933 (F), followed by determination of mitochondrial membrane potential of HSCs by FACS. The FACS plots with the gating for the effect of KU55933 in the KO mice are shown in (G) where individual mouse is represented by the individual line, and the results are quantified in (H), n=3 for each. Data are shown as mean ±SD for all bar graphs. *P<0.05, **P<0.01, ***P<0.001, by 1-factor ANOVA with post hoc *t* test. Scale bars, 400 µm. See also Figure S6.

We also attempted to rescue the function of the DPY30-deficient HSC in vivo by suppressing DDR. To this end, we have not been able to rescue the hematopoietic defects of DPY30-deficient mice by in vivo administration of ATM inhibitor based on peripheral blood profiling. However, the mitochondrial membrane potential, which was reduced by DPY30 loss, was consistently (though did not reach statistical significance) increased by DDR suppression in the DPY30-deficient mice, although such increase was mainly mediated by an apparently new population of cells among the phenotypic HSCs in these mice (**Figures 5F-H**). These results suggest a connection of the DDR, energy metabolism, and HSPC function, all under the control of the epigenetic regulator DPY30.

### Induction of *p21* is partially responsible for the functional defects of the DPY30-deficient HSPCs

Considering the established role of P21 in regulation of cell cycle and stem cell quiescence in response to metabolic and genotoxic stresses, we next sought to determine if the specific and dramatic elevation of *p21* expression played a role in HSC defect upon DPY30 loss. We generated *Dpy30*^*F/F*^; *p21*^*-/-*^; *Mx1-Cre* mice, which, upon pIpC injection, became *Dpy30* and *p21* double KO in the hematopoietic system. *p21* KO had no effect on the numbers or frequencies of any hematopoietic cell populations when DPY30 was present. However, upon *Dpy30* deletion, which caused severe accumulation of early HSPCs, *p21* KO significantly alleviated the aberrant accumulation of phenotypic LSK cells (**Figures 6A and S7A**).

**Figure 6.**
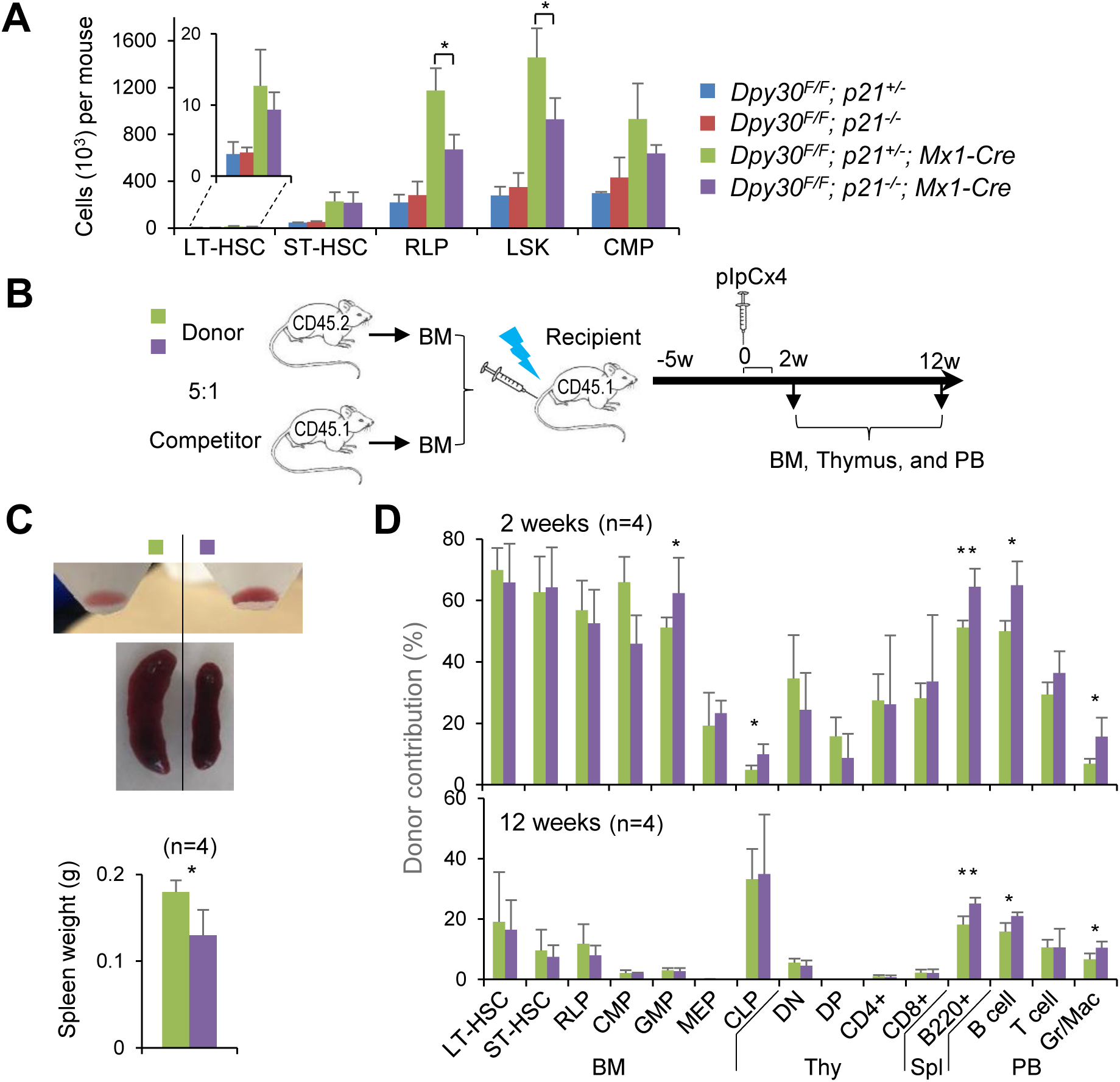
*P21* deletion partially rescues the function of the *Dpy30* KO hematopoietic cells. **(A)** Absolute numbers of BM cell populations in pIpC-injected mice of indicated genotypes. n=3 each. **(B)** Scheme for the mixed chimera transplantation system using whole BM cells from *Dpy30*^*F/F*^; *p21*^*+/-*^; *Mx1-Cre* or *Dpy30*^*F/F*^; *p21*^*-/-*^; *Mx1-Cre* mice as donors and whole BM cells from wild type mice as competitors. **(C)** Representative whole BMs flushed from long bones (top) and spleens (middle) in recipient mice (that received indicated donor cells) at two weeks after pIpC injection. Spleen weights are quantified (bottom). n=4 for each. **(D)** Contribution of indicated donors to different cell populations in BM chimeras at two and 12 weeks after pIpC injection. n=4 for each. Data are shown as mean ±SD for all bar graphs. *P<0.05, **P<0.01, by 1-factor ANOVA with post hoc *t* test (A) and 2-tailed Student’s t-test (C) and (D).. See also Figure S7.

We then transplanted donor BM from the *Dpy30*^*F/F*^; *p21*^*-/-*^; *Mx1-Cre* and *Dpy30*^*F/F*^; *p21*^*+/-*^; *Mx1-Cre* mice into irradiated recipient mice (**Figure 6B**). Two weeks after pIpC injection, we found that the recipients transplanted with the *Dpy30*^*F/F*^; *p21*^*-/-*^; *Mx1-Cre* BM had noticeably more red blood cells than those with the *Dpy30*^*F/F*^; *p21*^*+/-*^; *Mx1-Cre* BM (**Figure 6C**). As the hematopoietic system of the irradiated recipient mice was mainly contributed by the donor BM, this result indicates that the hematopoietic functionality damaged by DPY30 deficiency was partially rescued by the compound deficiency of P21. Consistent with the improved red blood cell generation, the recipients with double KO donor BM had significantly smaller spleen than those with *Dpy30*-KO donor BM (**Figure 6C**), suggesting that P21 deficiency alleviated splenomegaly induced by DPY30 loss. Moreover, at 2 weeks after *Dpy30* deletion, the double KO donors contributed significantly more than the *Dpy30* KO donors to a subset of hematopoietic cell populations, including GMP and CLP cells in the BM, B cells in spleen, and B cells and Gr/Mac cells in the periphery blood. At 12 weeks after *Dpy30* deletion, this effect was sustained for the splenic and periphery blood cell populations (**Figures 6D and S7B**). These results support a functional role of P21 in mediating the regulation of HSPCs by DPY30.

## DISCUSSION

Together with our previous results (Yang et al., 2016), the data in this work support a model for the role of DPY30 in fetal and adult HSPCs through regulation of energy metabolism and genome integrity (**Figure 7**). In normal HSPCs, DPY30, via its activity in facilitating H3K4 methylation, ensures the HSPC functionality through multiple pathways. These pathways include expression of key regulatory genes for adult HSC maintenance (Yang et al., 2016) and likely a different set of regulatory genes for fetal HSC maintenance, expression of key metabolic genes to maintain appropriate bioenergetic pathways, and safeguarding genome integrity through promoting efficient DNA repair. In the absence of DPY30 and efficient H3K4 methylation, expression of genes in many of these pathways is dysregulated, leading to dysregulation of energy metabolism and increase in DNA damage. The impairment in glycolysis may affect HSC maintenance, and the impairment in mitochondrial function may disable HSC activation. These stresses induce responses including P21 and other inflammatory pathways, which will suppress the activity of HSPCs and lead to attrition of stem cells in the long term. Considering that loss of SETD1A, one of the catalytic subunits responsible for the bulk level of cellular H3K4 methylation, increases the sensitivity of HSCs to inflammatory challenges (Arndt et al., 2018), we cannot formally exclude the potential complications by the inflammation-driven *Dpy30* ablation by the Mx1-Cre system. However, our conclusions were supported by alternative methods of *Dpy30* ablation in this and previous studies (Yang et al., 2016).

**Figure 7.**
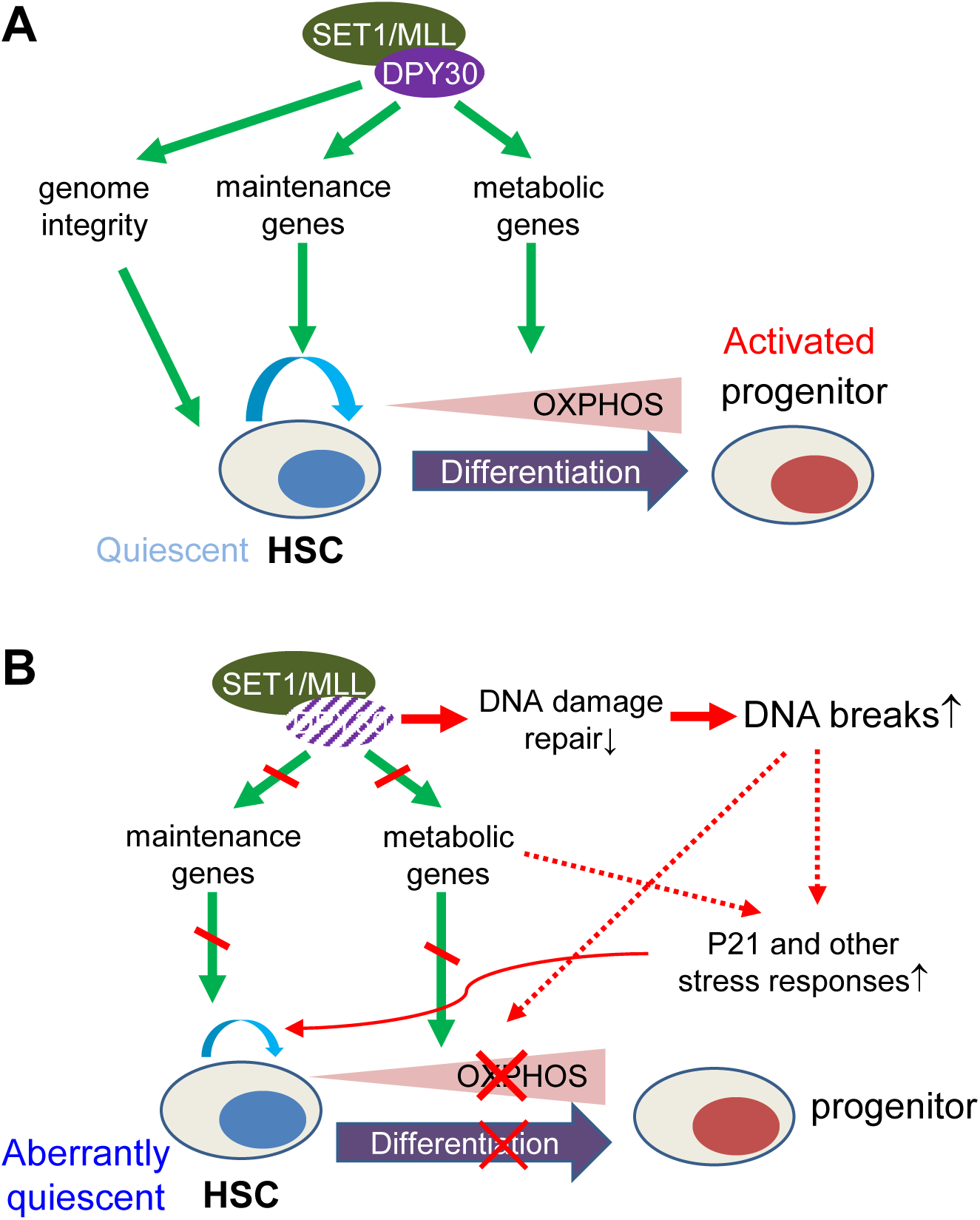
A model for the role of DPY30 in HSPC through regulation of energy metabolism and genome integrity. **(A)** DPY30 normally ensures the functionality of HSPC through guarding genome integrity and regulating key maintenance genes. It also enables HSC activation by regulating genes in energy metabolism. **(B)** DPY30 loss results in dysregulation of metabolism and key maintenance gene expression, as well as breach of genome integrity. These stresses induce P21 and other responses that lead to stem cell loss and activation defect.

We note that our gene analyses for BM used cells from donor-derived cells in the transplant recipients (Yang et al., 2016), while those for FL here used cells in a homeostatic setting without transplant. As a result, the commonly affected genes by DPY30 deficiency in both FL and BM HSCs identified in this work are likely underestimated, but represent genes truly responsive to DPY30 loss regardless of developmental stage or homeostatic/transplant settings. These genes are likely important in mediating the effect of DPY30, but may alternatively reflect a compensatory effect following DPY30 loss and alteration of certain important biological pathways. The common upregulation of a number of enzymes in oxidative metabolism in both FL and BM HSCs following DPY30 deficiency is likely a compensatory response to the impaired mitochondrial function. The overall level of ATP was not significantly perturbed in DPY30-deficient HSCs, likely due to compensatory responses from other pathways, such as potential increase in fatty acid catabolism reflected by the increased phosphorylation (**Figure 2A**) and inhibition of its negative regulator, ACC (Viollet et al., 2009). However, the unaffected ATP level makes it unlikely that the HSC dysfunction is a result of reduced total cellular energy level. Rather, we favor the possibility that the altered usage of bioenergetics pathways affects the HSC functionality. DPY30 deficiency appears to lead to a deeper quiescence of the HSCs based on the cell cycle entry. As quiescent HSCs are known to rely on glycolysis but increase oxidative phosphorylation and energy production upon activation (Kohli and Passegue, 2014; Yu et al., 2013), our results suggest that DPY30 plays important role in enabling HSC activation by ensuring metabolic reprogramming at the chromatin level.

A number of mouse models with deleted chromatin modulators (Liu et al., 2009; Santos et al., 2014; Tasdogan et al., 2016) and consequent HSPC defects exhibit increase in DDR and ROS level, and the excessive ROS level was shown to be a major contributor to the altered HSPC activity. Consistent with an increase in ROS upon depletion of DPY30 in human fibroblast cells (Simboeck et al., 2013), our results here demonstrate that the ROS level was increased in differentiated adult cells. However, we found a marked reduction of ROS in early HSPCs in the adult stage and in all hematopoietic cells in the prenatal stage upon depletion of DPY30 and H3K4 methylation, consistent with the reduced mitochondrial activity. Loss of the Setd1a also leads to decrease in ROS in HSCs, although the effect on mitochondrial function was not examined (Arndt et al., 2018). Our data show that the differential increase of ROS in lineage-committed cells functionally contributes to the reduced erythropoiesis in the DPY30-deficient animals as it can be partially rescued by antioxidant treatment, but we do not know the functional impact of the abnormally low level of ROS on HSPC activity. It appeared that the DPY30-deficient HSCs in both FL and BM made an effort to increase ROS level by upregulating expression of ROS-producing enzymes, suggesting a need to maintain a certain level of ROS for normal HSC function. This is consistent with previous findings that moderate levels of intracellular ROS are needed to activate DNA repair pathway and maintain genomic stability in stem cells (Li and Marban, 2010), and also to support HSC proliferation, differentiation, and mobilization (Chaudhari et al., 2014).

The important function of DPY30 and H3K4 methylation in efficient DNA repair is likely mediated through two non-exclusive mechanisms. First, DPY30 and H3K4 methylation are important for appropriate expression of certain DNA repair genes. This echoes the finding that SETD1A loss also impairs DNA repair capacity and results in increased DNA breaks, probably due to reduced H3K4 methylation and expression of multiple DNA repair genes (Arndt et al., 2018). Interestingly, efficient DDR gene expression in leukemia cells requires not the H3K4 methylation activity, but a novel region of SETD1A that binds to CYCLIN K, and H3K4 methylation at the DDR genes was sustained by other methylases in the absence of SETD1A in leukemia cells (Hoshii et al., 2018). Second, H3K4 methylation may directly regulate DNA repair through impact on the chromatin setting at the damaged sites. H3K4 methylation plays an important role in efficient DNA repair in yeast (Faucher and Wellinger, 2010) and *C. elegans* (Herbette et al., 2017), but may also predispose chromatin for DNA double stranded breaks through decondensing local chromatin (Burman et al., 2015). As DPY30-associated H3K4 methylation is functionally important for chromatin accessibility (Yang et al., 2018), it may regulate the efficiency of DNA repair by promoting the accessibility of DNA repair proteins. In addition to facilitating DNA repair, H3K4 methylation may also protect nascent DNA from degradation in replication stress (Higgs et al., 2018), thereby contributing to the maintenance of genome stability. Our results also suggest that it is the response to DNA damage, rather than the damage itself, that results in the inactivity of HSPCs. While suppression of DDR can temporarily rescue the function of HSPCs, we speculate that it would have a long-term consequence on the system with uncleansed HSPCs containing damaged genetic information.

In line with P21 being important in maintaining HSC quiescence (Cheng et al., 2000), its level is uniquely (among all CDK inhibitors) and greatly elevated in the DPY30-deficient HSCs in both FL and BM, keeping the cells in aberrantly deep quiescence regarding the cell cycle entry. Removing this quiescence maintenance factor helps release the DPY30-deficient HSCs from the deep quiescence and partially enables them to be activated and contribute to the hematopoietic reconstitution after transplant. Similar to the DDR inhibition, we speculate that such release would be mutagenic in the long term of the animals’ life span. In another study (Shah et al.), we have shown upregulation of *p21* and *p57* (*Cdkn1c*) upon DPY30 loss in the neural stem cell-enriched brain regions, suggesting that upregulation of CDK inhibitors as a general pathway in limiting the activity of tissue-specific stem and progenitor cells upon impaired epigenetic modifications following DPY30 loss. The incomplete rescue by *p21* KO suggests that the metabolic and genotoxic stresses upon DPY30 loss may lead to HSPC dysfunction through other effectors. For example, mitochondrial inactivation can lead to HSC defects through metabolite imbalances and epigenetic dysregulation (Anso et al., 2017).

Taken together, our results demonstrate a profound control of HSPC fate determination by a key chromatin modulator via regulating energy metabolism and genome integrity. The functional interplay among the metabolic regulation, ROS level, DDR, and the HSPC activities warrants further studies. Considering a critical role of DPY30 in *MLL*-rearranged leukemogenesis (Yang et al., 2014) and MYC-driven lymphomagenesis (Yang et al., 2018), it will be of great interest to investigate whether and how inhibition of this key epigenetic modulator affects cellular metabolism and genome integrity as part of mechanisms underlying cancer suppression.

## EXPERIMENTAL PROCEDURES

### Animals

All animal procedures were approved by the Institutional Animal Care and Use Committee at the University of Alabama at Birmingham. All mice were maintained under specific pathogen free conditions and housed in individually ventilated cages. *Dpy30*^*+/-*^ mice were generated in our laboratory as previously reported (Yang et al., 2016) and were crossed to *Mx1-Cre* (Jackson Laboratory, JAX 003556), and *Vav-Cre* (Jackson Laboratory, JAX 008610) mice to produce *Cre; Dpy30*^*+/-*^ Mice. These mice were further crossed with *Dpy30*^*F/F*^ mice to produce *Cre; Dpy30*^*F/+*^ and *Cre; Dpy30*^*F/-*^ littermates for experimental use. All transplant recipient mice were C57Bl/6J and CD45.1^+^, and purchased from Charles River Laboratories. *P21*^*-/-*^ (*129S2-Cdkn1atm1Tyj/J*) mice (Jackson Laboratory, JAX 003263) were further crossed with *Mx1-Cre* and *Dpy30*^*F/F*^ to generate the littermates of *Dpy30*^*F/F*^; *P21*^*+/-*^; *Mx1-Cre* and *Dpy30*^*F/F*^; *P21*^*-/-*^; *Mx1-Cre*.

### Measurement of oxygen consumption, NAD^+^/NADH, NADP^+^/NADPH, and total ATP level

Measurement of intact cellular respiration was performed using the Seahorse XF24 analyzers. Briefly, the respiration of BM Lin^-^ cells was measured under basal conditions, in the presence of mitochondrial inhibitor oligomycin (500 nM), mitochondrial uncoupling compound carbonylcyanide - 4-trifluoromethoxyphenylhydrazone (FCCP) (4 µM), and respiratory chain inhibitor rotenone (1 µM). Sorted BM Lin^-^ Sca1^+^cKit^+^ (LSK) cells were suspended in PBS and 10,000 cells were added to each well. The ratios of NAD^+^/NADH, NADP/NADPH and GSH/GSSG were measured with NAD^+^/NADH-Glo Assay, NADP/NADPH-Glo Assay and GSH/GSSG-Glo Assay Luminescent Cell Viability Assay kits (all from Promega, Madison, WI), respectively, according to manufacturer’s instructions. To determine total cellular ATP levels, BM LSK cells were sorted and assessed using a CellTiter-Glo Luminescent Cell Viability Assay kit (Promega Corporation).

### Flow cytometry for analysis and cell isolation

Single cell suspensions were prepared from FL, BM, thymus, spleen, or peripheral blood and stained with antibodies as previously described (Yang et al., 2016).

### Statistics

Unless indicated in the figure legends, the unpaired 2-tailed Student’s t-test was used to calculate P values and evaluate the statistical significance of the difference between indicated samples. A P value less than 0.05 was considered significant. Four groups comparison was analyzed by one-factor ANOVA as indicated in figure legends. If ANOVA was overall significant, post hoc *t* test was used for pairwise comparisons of interest.

### Data availability

The RNA-seq data have been deposited in Gene Expression Omnibus database with the accession number GSE101856.

## AUTHOR CONTRIBUTIONS

Z.Y. and H.J. conceived the project and designed the experiments. Z.Y. conducted most of the experiments and analyzed the results, K.S. conducted some of the experiments and analyzed the results, A.K. conducted bioinformatic analyses. Z.Y. and H.J. wrote the paper.

## ACKNOWLEDGEMENTS

We thank the Comprehensive Flow Cytometry Core (CFCC) at UAB, which are supported by NIH core grants P30 AR048311 and P30 AI027767. The X-RAD 320 irradiator was purchased by the UAB animal facility using NIH Grant G20RR022807. We thank Ying Gai Tusing and Yanfang Zhao for technical assistance with mice work. This work was supported by NIH grant R01DK105531, Start-up funds from the State of Alabama and University of Virginia. HJ is a recipient of the American Society of Hematology Scholar Award, the American Cancer Society Research Scholar Award, and the Leukemia and Lymphoma Society Scholar Award.

## SUPPLEMENTAL INFORMATION

**Figure S1.**
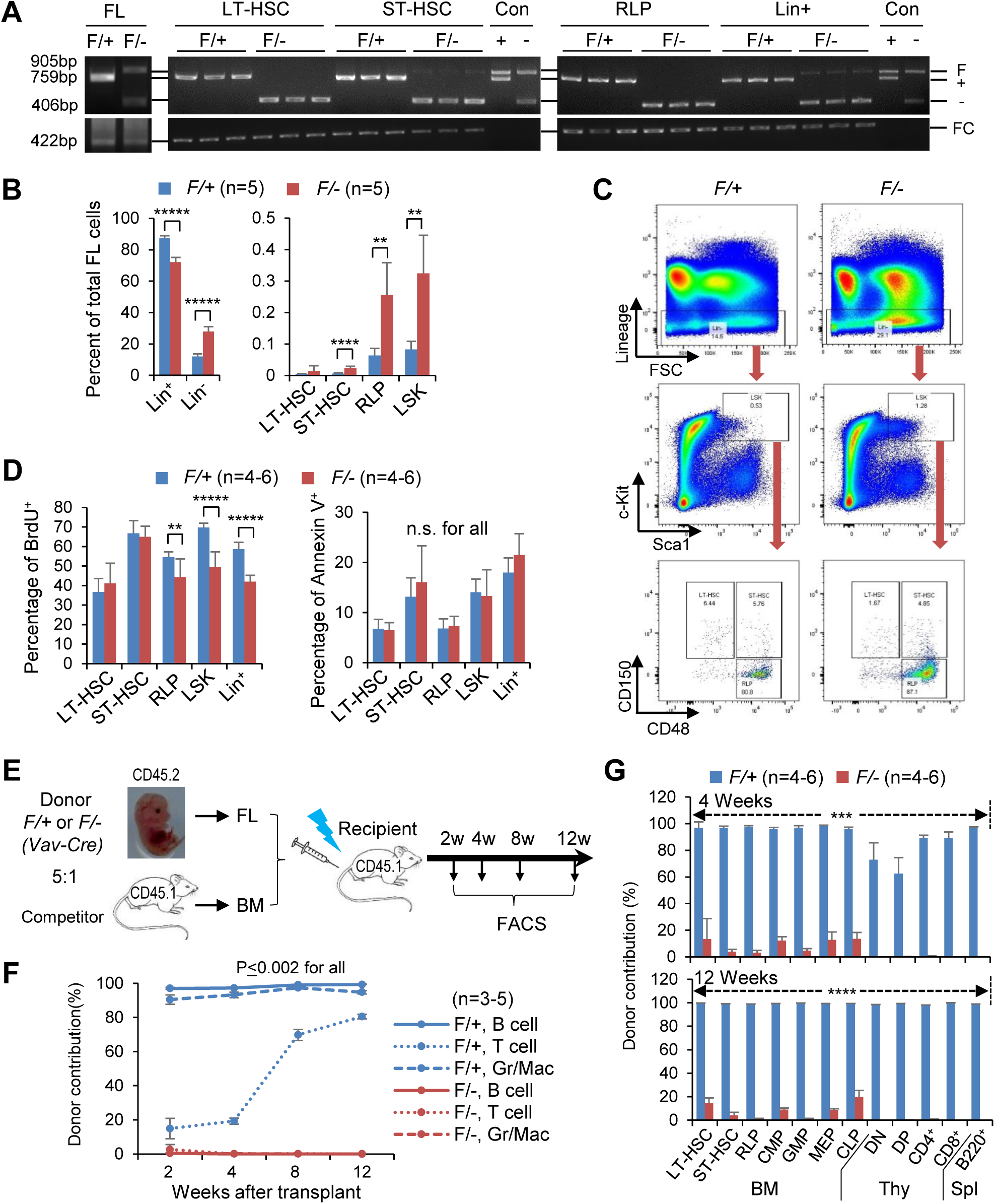
Dpy30 deficiency in fetal liver results in anemia and defective HSC function. *Vav-Cre; Dpy30*^*F/+*^ (*F*/+) and *Vav-Cre; Dpy30*^*F/-*^ (*F*/-) littermate fetuses were used in all panels. **(A)** Genomic PCR to detect *Dpy30* deletion in total fetal liver and FACS sorted fetal liver cell populations as indicated at E14.5. Each lane was from an embryo. The calculated sizes of the PCR products (shown on the right) are consistent with the mobility of the bands. Note that the recombination in Lin+ cells was inefficient. Con (+) and Con (-) were from *Dpy30*^*F/+*^ and *Dpy30*^*F/-*^ (no Cre) fetal liver cells, respectively. **(B)** Percentage of different cell populations in fetal liver. n=5 each. **(C)** Representative FACS analysis of fetal liver cells at E14.5. **(D)** BrdU incorporation (left) and annexin V staining (right) assays for different cell populations in fetal liver at E14.5. n=4-6. **(E)** Scheme for the competitive transplantation system using whole fetal liver cells from *Vav-Cre*; *Dpy30*^*F/+*^ and *Vav-Cre*; *Dpy30*^*F/-*^ as donors and wild type BM as competitors. **(F)** Donor contribution to indicated peripheral blood cells was determined by FACS at different times post transplantation following the scheme in (e). n=3-5 mice. **(G)** Donor contribution to different cell populations in bone marrow (BM), thymus (Thy) and spleen (Spl) at 4 weeks (top) and 12 weeks (bottom) after transplantation. n=4-6 mice. Data are shown as mean ± SD for all bar graphs and (F). n.s., not significant, **P<0.01, ***P<0.001, ****P<0.0001, ***** P<0.00001, 2-tailed Student’s *t*-test. Related to Figure 1.

**Figure S2.**
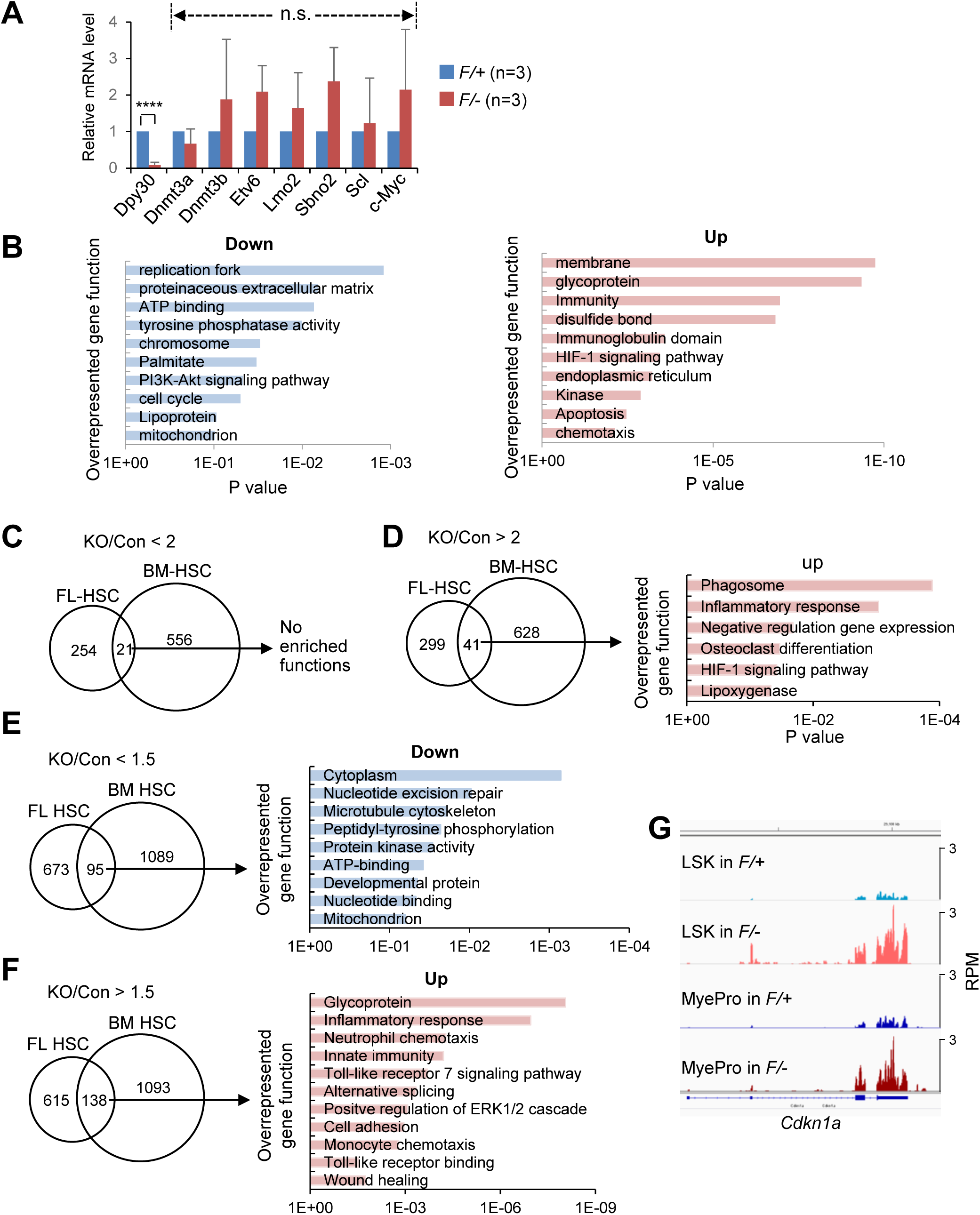
Analysis of genes significantly altered by Dpy30 loss in HSCs. **(A)** Relative mRNA levels of indicated genes in the HSCs sorted from fetal livers of *Vav-Cre; Dpy30*^*F/+*^ and *Vav-Cre; Dpy30*^*F/-*^ littermate fetuses at E14.5 were determined by RT-qPCR and normalized to *Actb*. Data are shown as mean ± SD. n=3 each. n.s., not significant, **** P<0.0001, by 2-tailed Student’s *t*-test. **(B)** Gene ontology analysis by DAVID for genes that were downregulated or upregulated over 2 fold in HSCs from 3 pairs of fetal littermates. **(C and D)** Venn diagram for genes that were downregulated (C) or upregulated (D) over 2 fold in HSCs from 3 pairs of fetal littermates and 2 pairs of BM HSCs (left). Gene ontology analysis by DAVID of the shared genes are shown (right). In panels (C-G), the BM HSC data are based on RNA-seq results in donor (*Mx1-Cre; Dpy30*^*F/+*^ or *Mx1-Cre; Dpy30*^*F/-*^) - derived HSCs in the BM chimera recipients in two independent BM transplantations (TP1 and TP2) from our previous work (Yang et al., 2016). **(E and F)** Venn diagram for genes that were downregulated (E) or upregulated (F) over 1.5 fold in HSCs from 3 pairs of fetal littermates and 2 pairs of BM HSCs (left). Gene ontology analysis by DAVID of the shared genes are shown (right). **(G)** RNA-seq profiles at *Cdkn1a* (*p21*) from one BM chimera transplant assay. Related to Figure 1.

**Figure S3.**
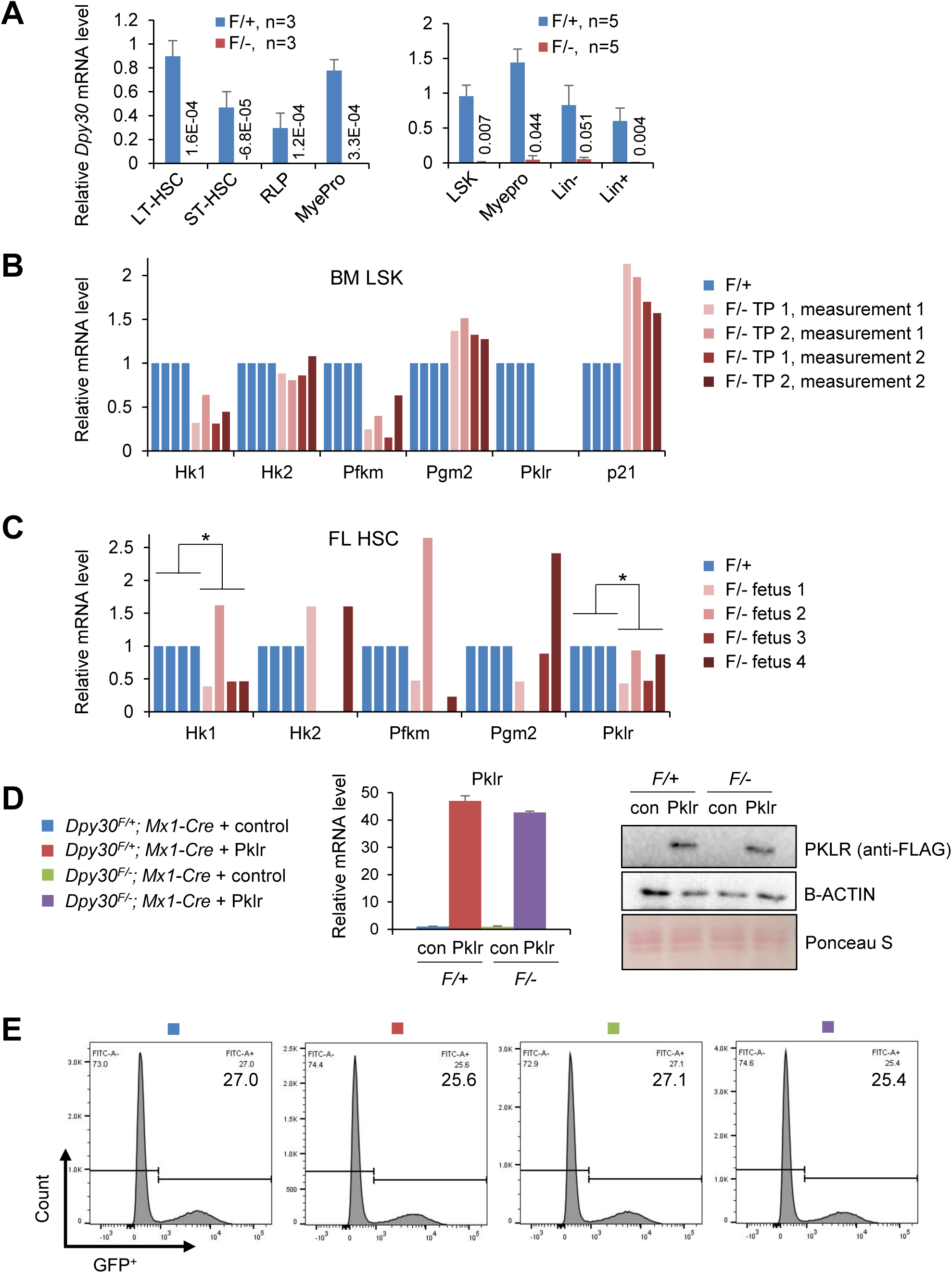
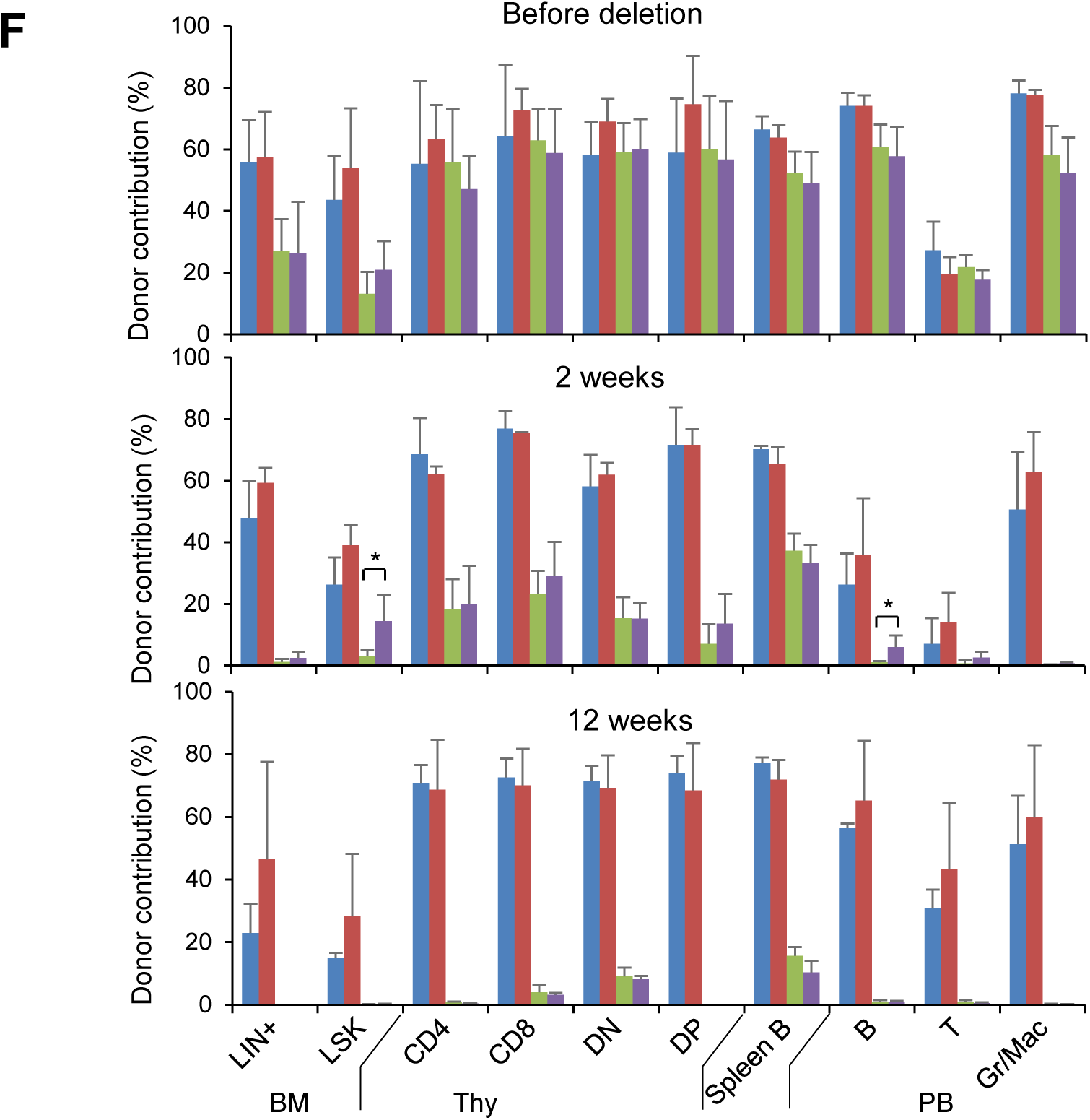
Downregulation of key glycolytic genes by Dpy30 loss. **(A)** Relative mRNA levels of *Dpy30* by RT-qPCR and normalized to *Actb*, in FACS sorted BM cell populations from *Mx1-Cre; Dpy30*^*F/+*^ (*F/+*) and *Mx1-Cre; Dpy30*^*F/-*^ (*F/-*) mice after pIpC injections. **(B and C)** Relative mRNA levels of indicated genes in the donor (*Mx1-Cre; Dpy30*^*F/+*^ or *Mx1-Cre; Dpy30*^*F/-*^) - derived LSK cells in the BM chimera recipients in two independent BM transplantations (TP1 and TP2) from previous work (Yang et al., 2016) for (B), and from fetal livers of *Vav-Cre; Dpy30*^*F/+*^ and *Vav-Cre; Dpy30*^*F/-*^ littermate fetuses at E14.5 for (C), by RT-qPCR and normalized to *Actb*, and the data are shown for each measurement. **(D)** Relative expression of FLAG-*Pklr* from retrovirally-transduced BM cells determined by RT-qPCR (left) and immunoblotting (right) prior to the transplant. The *Pklr* mRNA levels were normalized to *Actb* and relative to the level in the *Dpy30*^*F/+*^; *Mx1-Cre* + control, and are shown as mean ±SD of technical duplicate. Anti-FLAG antibody was used as our anti-PKLR antibody failed to react with mouse PKLR. Legends are the same in (D-F). **(E)** FACS analysis of the retroviral transduction efficiency of BM cells, shown by the percentage of GFP-positive population. **(F)** Donor contribution to indicated cell populations in bone marrow (BM), thymus (Thy), spleen, and peripheral blood (PB) before deletion (top), 2 weeks (middle) and 12 weeks (bottom) after pIpC-injections in the transplant recipients following scheme in Figure 2G. n=4 each. Data are shown as mean ±SD. *P<0.05, **P<0.01, ***P<0.001, by 2-tailed Student’s *t*-test. Related to Figure 2.

**Figure S4.**
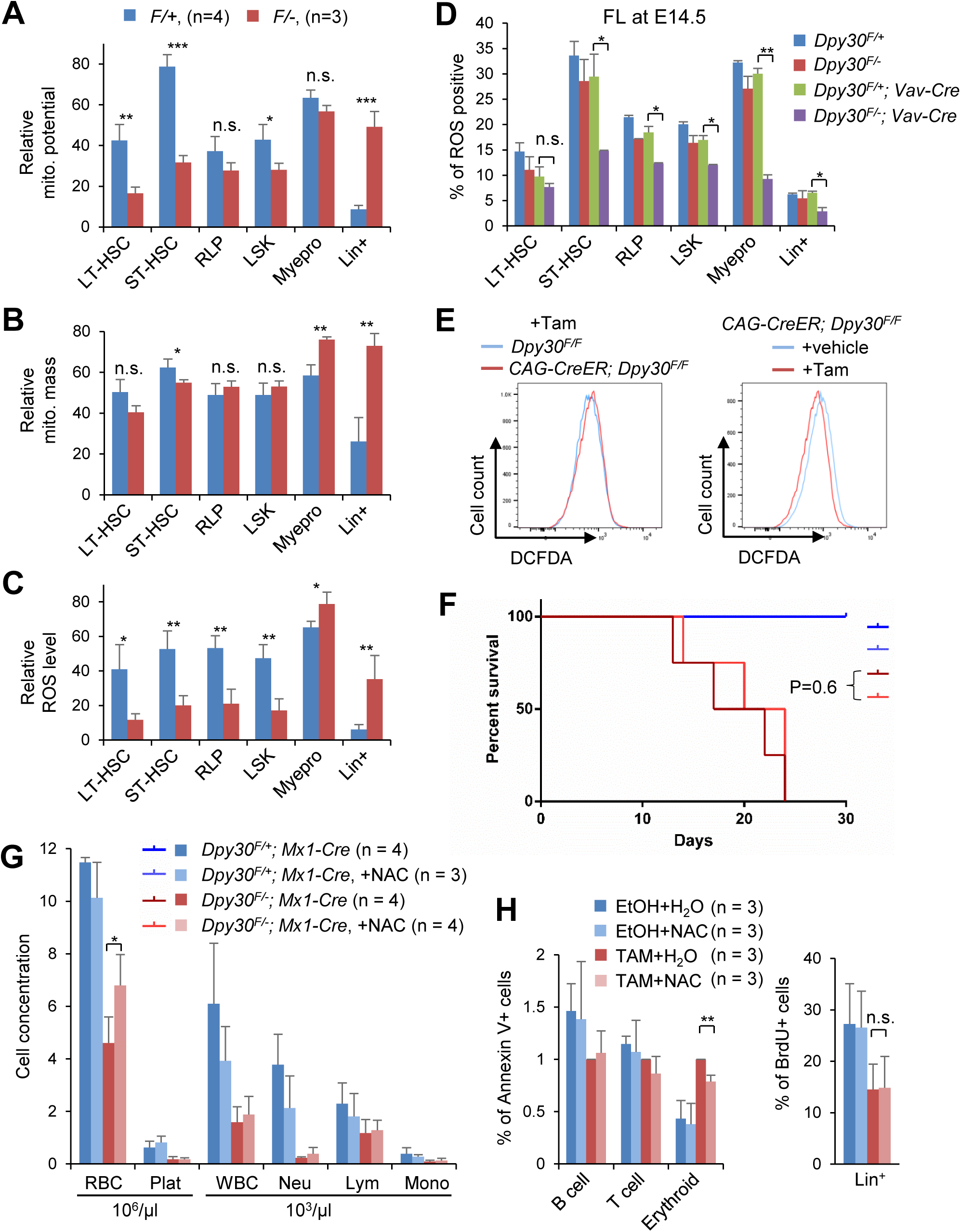
Dpy30 deficiency impairs mitochondrial function in different types of cells. **(A-C)** Mitochondrial membrane potentials (A), mitochondrial mass (B) and intracellular ROS levels (C) in different BM cell populations from pIpC-injected *Mx1-Cre; Dpy30*^*F/+*^ (*F/+*) and *Mx1-Cre; Dpy30*^*F/-*^ (*F/-*) mice were determined by FACS. Because these values were vastly different for MyePro and Lin^+^ cells (much higher) compared to the cell populations, gating criterion for these values were the same for LT-HSC, ST-HSC, RLP, and LSK cells, but different for MyePro, and that for Lin^+^ cells was also different from all other cells. **(D)** Intracellular ROS levels in different cell populations of fetal livers of indicated genotypes at E14.5. n=2 each. **(E)** Primary MEFs of indicated genotypes were treated with vehicle (ethanol) or 4-OH tamoxifen (Tam), and intracellular ROS levels were determined by FACS. **(F and G)** Effects of antioxidant administration on hematopoiesis of Dpy30-deficient animals. Legends in (G) apply to (F) and (G). Indicated number of pIpC-injected *Mx1-Cre; Dpy30*^*F/+*^ and *Mx1-Cre; Dpy30*^*F/-*^ mice were on NAC-containing drinking water or not, and were examined for survival (F) and peripheral blood profiling (G), **(H)** Lin^+^ BM cells from three *CAG-CreER; Dpy30*^*F/F*^ mice were treated with either EtOH control or tamoxifen (TAM), and water control or NAC in culture for two days, and subjected to assays for apoptosis by Annexin V staining and for cell proliferation by in vitro BrdU incorporation. Data are shown as mean ±SD for all panels. Number of animals are indicated. n.s., not significant, *P<0.05, **P<0.01, ***P<0.001, by 2-tailed Student’s *t*-test, except by 1-factor ANOVA with post hoc *t* test for (D) and log-rank test for (F). Related to Figure 3.

**Figure S5.**
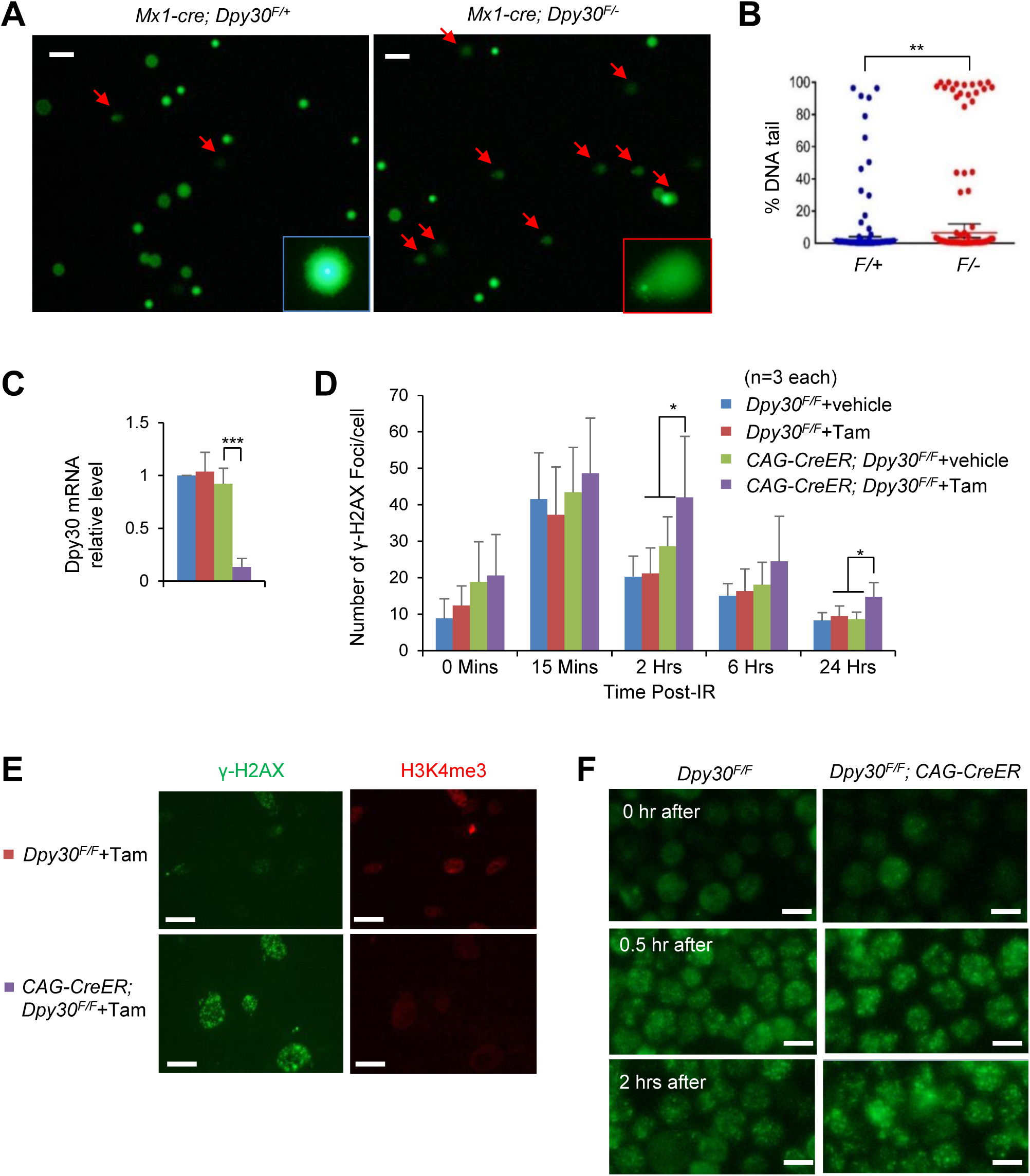
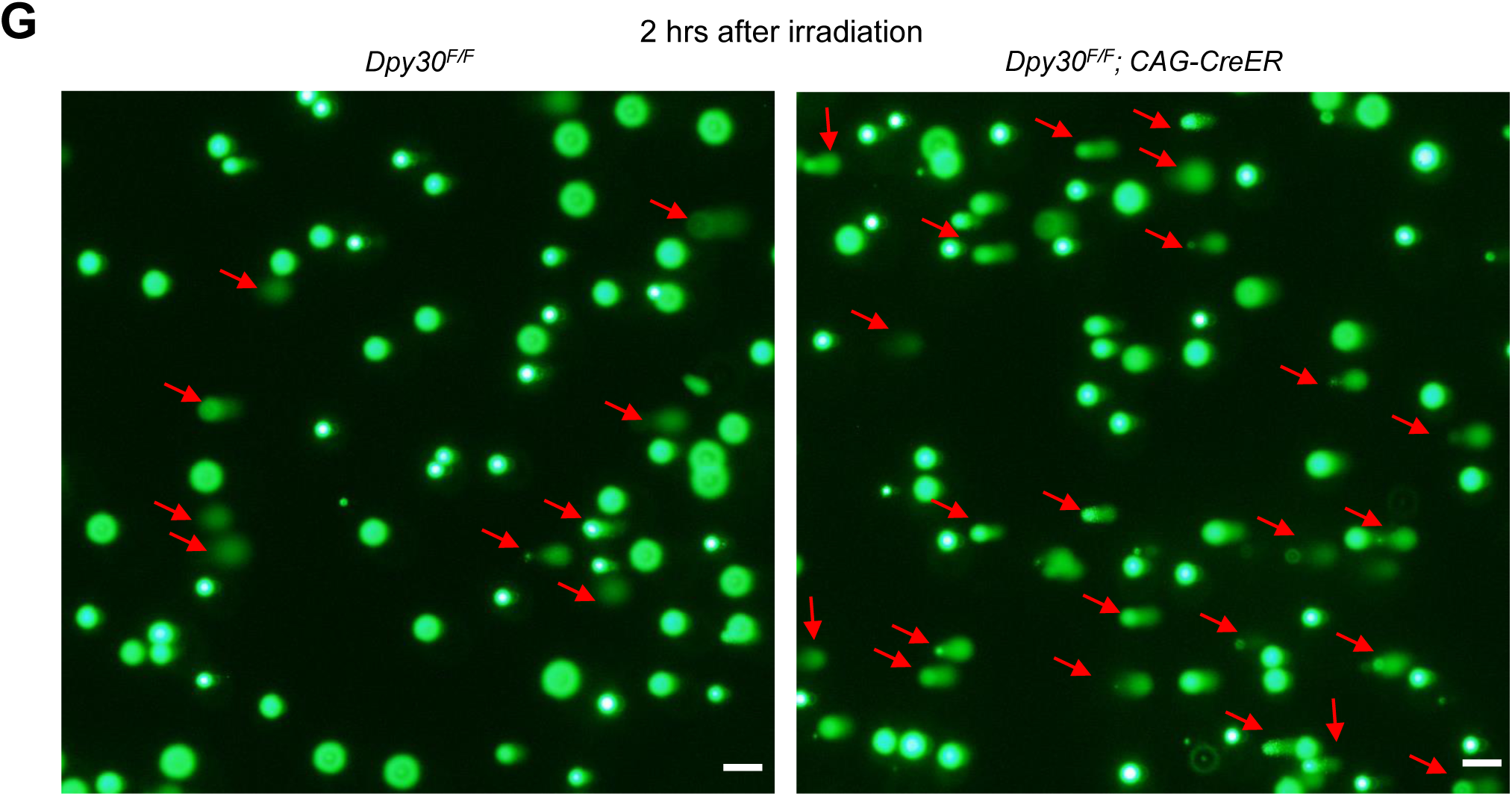
Dpy30 deficiency results in increase in DNA damage and impairs DNA repair. **(A and B)** Comet assay using BM LSK cells pooled from 3 mice each, with representative images in (A) and quantification of the relative tailing level of cells in (B). Red arrows point to comet-shaped cells with significant DNA damage. The representative zoom-in images of an undamaged (blue box) and damaged (red box) cells are shown in the insets. In (B), each dot represents a cell analyzed by the OpenComet software, and 100% means maximum tail DNA (most damage) and 0% means tail DNA (no damage). **(C-E)** Primary MEFs of indicated genotypes were treated with vehicle (ethanol) or 4-OH tamoxifen, and subjected to ionizing radiation at 0 min, followed by γ-H2AX staining at indicated times after radiation. Relative *Dpy30* mRNA levels were determined by RT-qPCR and normalized to *Actb*, n=2 for each of the left two bars and n=4 for each of the right two bars (C). The γ-H2AX foci numbers per cell from 20-60 cells in each repeat of 3 biological repeats are plotted (D). Representative images of indicated cells at 2 hrs post irradiation are shown (E). **(F and G)** Representative images of γ-H2AX staining (F) and comet assay (G) for LSK cells from indicated animals at indicated times post irradiation following the scheme in Figure 4B. (G) is from 2 hrs after irradiation. Data are shown as mean ±SD. *P<0.05, **P<0.01, ***P<0.001, by 2-tailed Student’s *t*-test for (B) and 1-factor ANOVA with post hoc *t* test for (C) and (D). Scale bars, 50 µm (S5A), 10 µm (S5E and S5F), 20 µm (S5G). Related to Figure 4.

**Figure S6.**
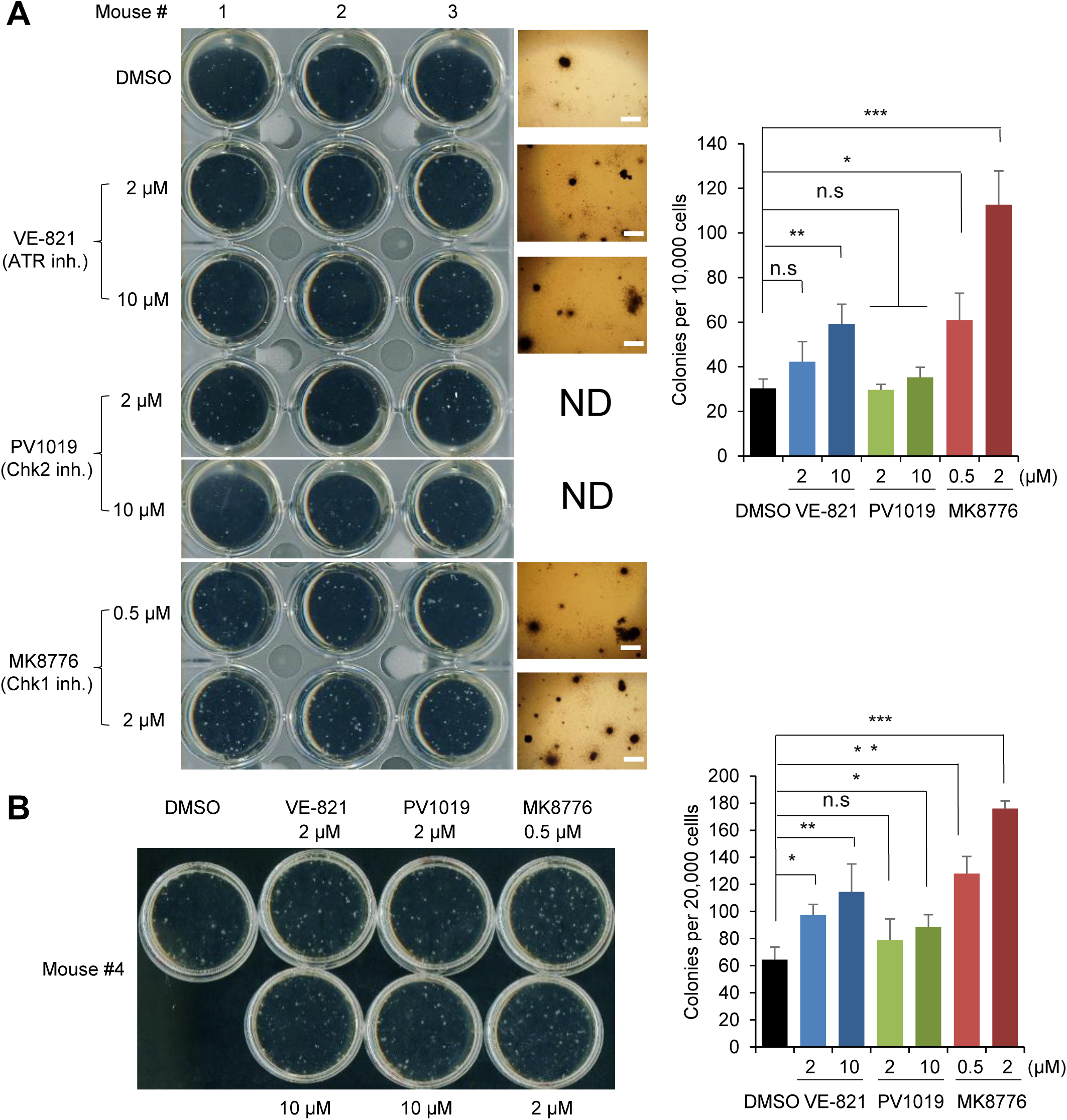
DDR Inhibition partially rescues the function of the Dpy30-deficient hematopoietic cells. Lin^-^ BM cells from *CAG-CreER; Dpy30*^*F/F*^ mice were treated with indicated agents in liquid culture for two days and subjected to colony formation assays in two independent sets of experiment in (A) and (B). In (A), 3 mice (#1, 2, 3) were used and 10,000 cells were seeded. In (B), 2 mice (#4, 5) were used and 20,000 cells were seeded. Representative image of the plates and colonies are shown on the left, and quantification for all colonies is shown on the right. ND, not done. Data are shown as mean ±SD for all bar graphs. *P<0.05, **P<0.01, ***P<0.001, by 1-factor ANOVA with post hoc *t* test. Scale bars, 300 µm. Related to Figure 5.

**Figure S7.**
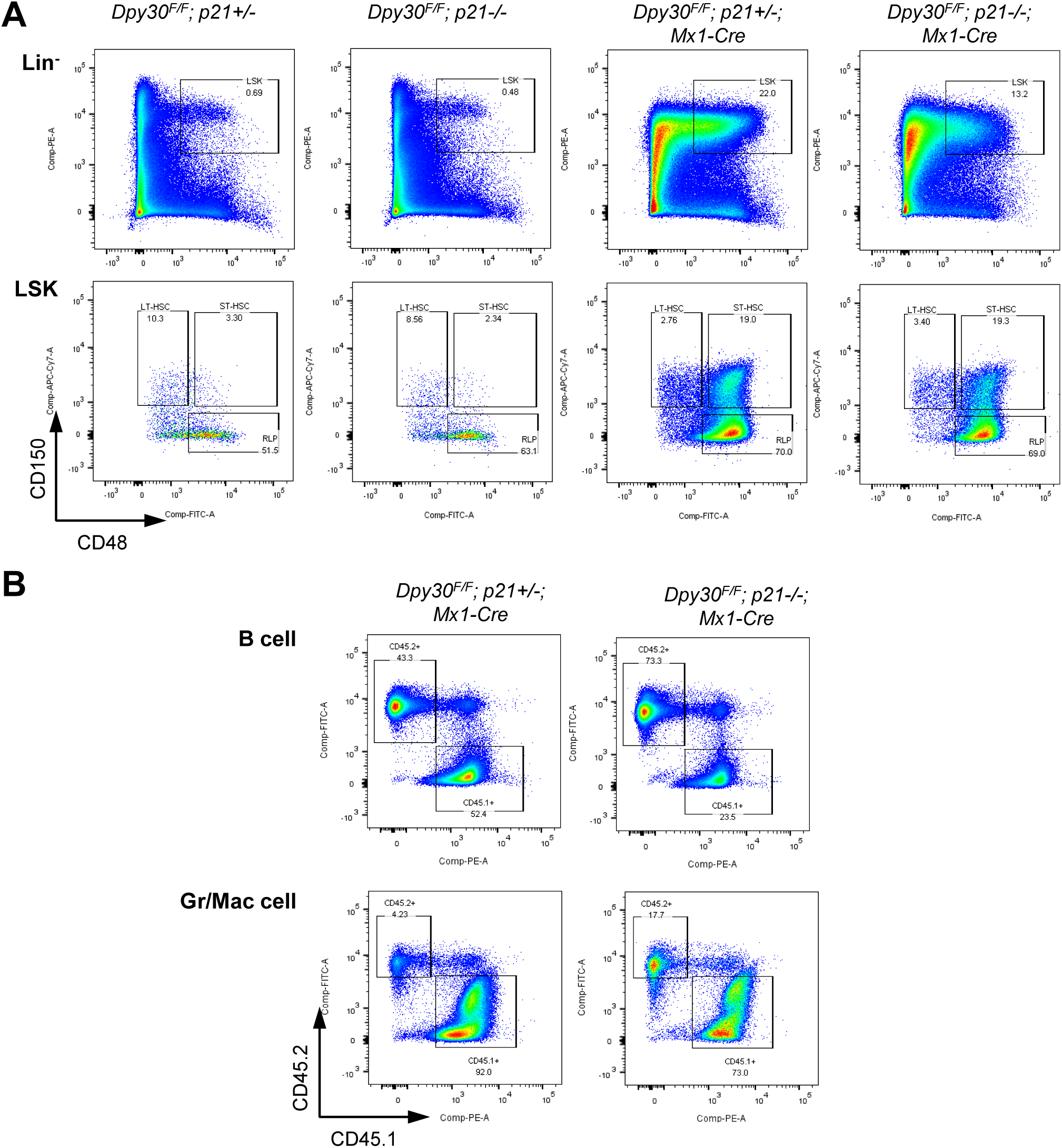
*P21* deletion partially rescues the function of the *Dpy30* KO hematopoietic cells. **(A)** Representative FACS analysis of BM from pIpC-injected mice of indicated genotypes. **(B)** Representative FACS analysis of B cells in spleen and granulocytes & macrophages in peripheral blood from the recipient mice that received donors of indicated genotypes (top) based on the scheme in Figure 6B. Related to Figure 6.

## SUPPLEMENTAL TABLES

**Table S1:** Cell surface phenotypes used in this work.

| Cell population | Cell surface phenotype |
| --- | --- |
| Lin<sup>-</sup> (Fetal liver) | CD3<sup>-</sup> CD4<sup>-</sup> CD5<sup>-</sup> CD8<sup>-</sup> B220<sup>-</sup> Gr1<sup>-</sup> TER119<sup>-</sup> |
| Lin<sup>-</sup> (Bone marrow) | CD3<sup>-</sup> B220<sup>-</sup> CD11b<sup>-</sup> Gr1<sup>-</sup> TER119<sup>-</sup> |
| Lin<sup>+</sup> | Cells excluding Lin<sup>-</sup> |
| LSK | Lin<sup>-</sup> Sca1<sup>+</sup> cKit<sup>+</sup> |
| LT-HSC | LSK CD48<sup>-</sup> CD150<sup>+</sup> |
| ST-HSC | LSK CD48<sup>+</sup> CD150<sup>+</sup> |
| RLP | LSK CD48<sup>+</sup> CD150<sup>-</sup> |
| MyePro | Lin<sup>-</sup> Sca1<sup>-</sup> cKit<sup>+</sup> (L<sup>-</sup> S<sup>-</sup> K<sup>+</sup>) |
| CLP | Lin<sup>-</sup> Sca1<sup>low</sup> cKit<sup>low</sup> IL7R<sup>+</sup> CD135<sup>+</sup> |
| CMP | Lin<sup>-</sup> Sca1<sup>-</sup> cKit<sup>+</sup> CD34<sup>+</sup> FcRγ<sup>-</sup> |
| MEP | Lin<sup>-</sup> Sca1<sup>-</sup> cKit<sup>+</sup> CD34<sup>-</sup> FcRγ<sup>-</sup> |
| GMP | Lin<sup>-</sup> Sca1<sup>-</sup> cKit<sup>+</sup> CD34<sup>+</sup> FcRγ<sup>+</sup> |
| B cell | B220<sup>+</sup> |
| T cell | CD3ε<sup>+</sup> |
| DN | CD4<sup>-</sup> CD8<sup>-</sup> |
| Erythroid | Ter-119<sup>+</sup> |
| DP | CD4<sup>+</sup> CD8<sup>+</sup> |
| Gr/Mac | Gr1<sup>+</sup> or Mac1<sup>+</sup> |

**Table S2. Gene expression analyses in fetal liver HSCs in control and *Dpy30* KO background.**

This Table is separately attached as an Excel file. There are 14 tabs in this file. Fetal liver (FL) HSCs from 3 pairs of *Vav-Cre; Dpy30*^*F/+*^ (control) and *Vav-Cre; Dpy30*^*F/-*^ (KO) fetuses, each pair from the same litter, at E14.5 were sorted out for RNA-Seq. Gene expression data for the bone marrow (BM) HSCs are from our previous work (Yang et al., 2016) and are included here for convenience of comparison. The BM data are from two transplant (TP1 and TP2) recipients at two weeks after pIpC induction. These TP recipients received *Mx1-Cre; Dpy30*^*F/+*^ (control) and *Mx1-Cre; Dpy30*^*F/-*^ (KO) donor BM cells and were then injected with pIpC. Compared to control, genes up- or down-regulated more than 2 (or 1.5) fold in both BM TPs are shown in tabs BM up>2 (or 1.5) and BM down>2 (or 1.5). All normalized FL HSC RNA-Seq data with three fetuses of each genotype are listed in tab All genes in FL HSC. Genes up- or down-regulated in FL HSCs with mean value more than 2 fold and p <0.05 are listed in tabs FL down >2 and FL up>2. Genes up- or down-regulated more than 1.5 fold in all three littermate pairs are listed in tabs FL down >1.5 and FL up>1.5. Tabs BM & FL down>2 (or 1.5) show overlapping genes between tabs BM down>2 (or 1.5) and FL down >2 (1.5). Tabs BM & FL up>2 (1.5) show overlapping genes between tabs BM up>2 and FL up>2.

**Table S3:**
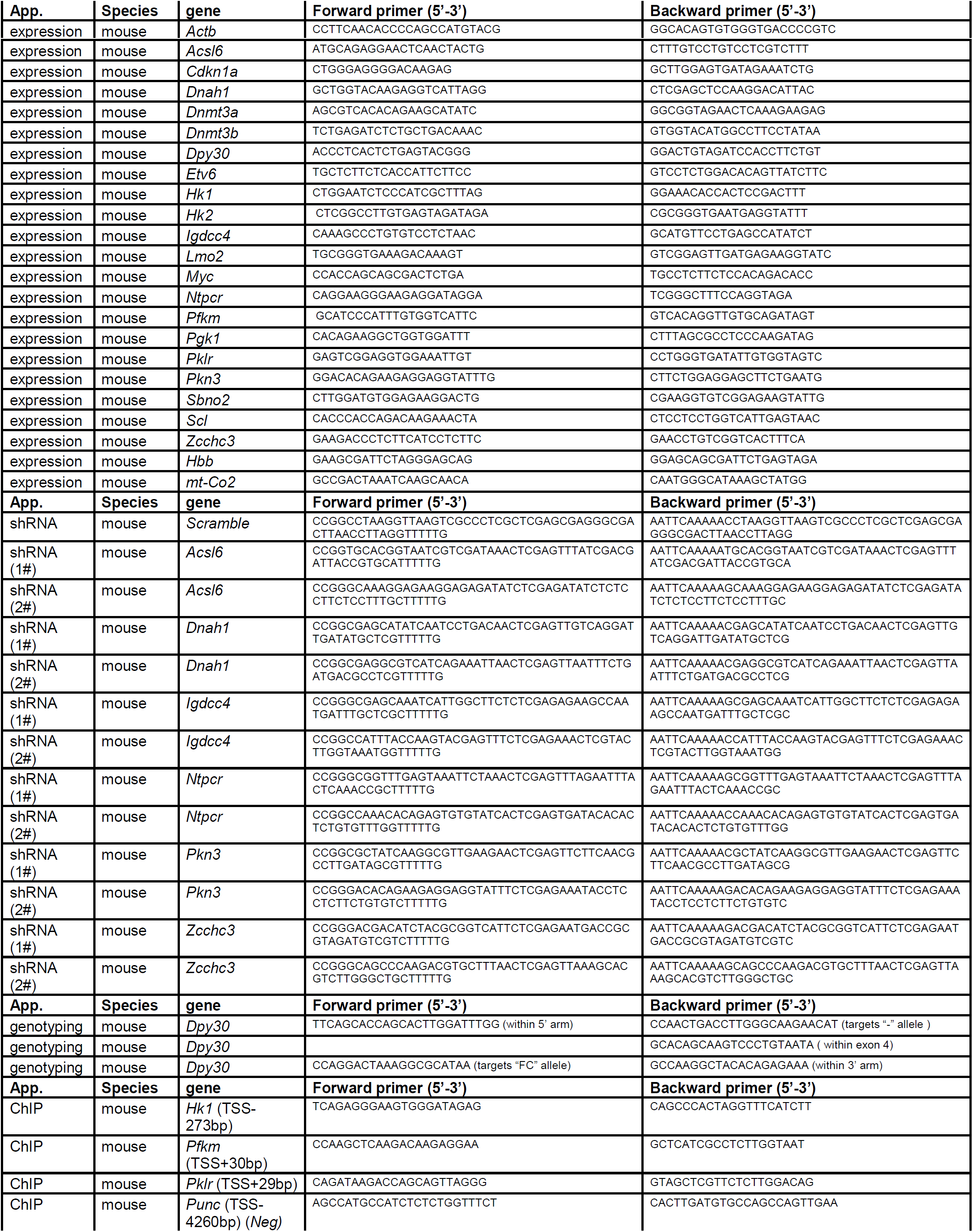
Primers and Oligos.

## SUPPLEMENTAL EXPERIMENTAL PROCEDURES

### Transplantation

For competitive transplantation assays, the donor FLs were separated from E14.5 embryos (CD45.2^+^, *Vav-Cre; Dpy30*^*F/+*^ and *Vav-Cre; Dpy30*^*F/-*^) and suspended in phosphate-buffered saline (PBS) supplemented with 3% heat-inactivated fetal bovine serum (FBS), followed by removal of red blood cells. 1 × 10^6^ donor FLs (CD45.2^+^) and 2 × 10^5^ whole BM cells (C57Bl/6J, CD45.1^+^) at a ratio of 5:1 were transplanted via tail vein into lethally irradiated recipient mice (CD45.1^+^). For mixed chimera assays, 1 × 10^6^ whole FLs as donors prepared from E14.5 embryos (CD45.2^+^, *Mx1-Cre; Dpy30*^*F/+*^ and *Mx1-Cre; Dpy30*^*F/-*^) were mixed with 2 × 10^5^ whole BM cells (C57Bl/6J, CD45.1^+^) at a ratio of 5:1, and transplanted into recipient mice in the same way as described above. Recipients were then injected with 150 µl pIpC (InvivoGen) at 1 mg/ml 4 times over 10 days starting at 5 weeks post transplantation. The first two injections were 2 days apart and last injections were 4 days apart. Reconstitution levels were assessed at different time points after pIpC injection. Prior to transplant, all of the recipient mice were lethally irradiated with a split dose of 1100 Rads with 3 hours apart. Donor contribution was determined by flow cytometry at different time points and calculated by CD45.2^+^/(CD45.1^+^ + CD45.2^+^) × 100% within the indicated cell type except for the Pklr rescue assay, where the donor contribution was calculated by GFP^+^/(GFP^+^ + CD45.1^+^) × 100%. For the Pklr rescue assay, the mouse *Pklr* cDNA was cloned with N-terminal FLAG tag into pMSCV-IRES-GFP (Addgene, #20672) between EcoRI and XhoI sites to generate the FLAG-*Pklr*-expressing plasmid. To generate the retroviruses, 293T cells were transiently transfected with either empty pMSCV-IRES-GFP vector or the FLAG-*Pklr*-expressing construct with pCL-Eco plasmid using Lipofectamine 2000 (Invitrogen), and viruses in the supernatants were harvested two days after transfection. Four days after injecting both *Dpy30*^*F/+*^; *Mx1-Cre* and *Dpy30*^*F/-*^; *Mx1-Cre* mice (6-8-week old) with 150 mg/kg 5-fluorouracil (5-FU, Sigma-Aldrich), BM cells were flushed from these mice and were incubated overnight in RPMI 1640 with 20% fetal calf serum, 20 ng/ml SCF, 20 ng/ml Flt-3, 10 ng/ml IL-3 and 10 ng/ml IL-6 (all from Peprotech, Rocky Hill, NJ) at 37°C to promote cell cycle entry. Cells were then spinoculated with retroviral supernatant in the presence of 8 μg/ml polybrene (Clontech, Mountain View, CA) for 90 minutes at 500 x g at 30°C. The spinoculated cells were analyzed for transduction efficiency (GFP positivity) by flow cytometry, and cells containing 1 × 10^5^ GFP^+^ cells (CD45.2^+^) and 2 × 10^5^ helper BM cells (CD45.1^+^) were transplanted via tail vein into lethally irradiated recipient mice (CD45.1^+^).

### Flow cytometry for analysis and cell isolation

Single cell suspensions were prepared from FL, BM, thymus, spleen, or peripheral blood and stained with antibodies as previously described (Yang et al., 2016). FACS analysis was performed on LSRFortessa (Becton Dickinson), and data were analyzed using FlowJo software (Tree Star) as previously described (Yang et al., 2016). Cells were fixed, permeablized and immunostained with anti-phospho-AMPKα (Thr172) antibody (Cell Signaling Technology, #2535), or phospho-KAP-1 (S824) (Bethyl Laboratories, #A300-767A). Mean fluorescence intensity was determined. To measure mitochondrial membrane potential and mitochondrial mass, the HSC staining was modified to make the channels available for tetramethyl rhodamine methyl ester (TMRM) and Mitotracker Deep Red staining (all from Molecular Probes), respectively. After antibody staining cells were incubated with 25 nM TMRM or 1nM Mitotracker Deep red for 15 min at 37°C followed by flow-cytometry on LSRFortessa (Becton Dickinson), respectively. To measure cellular ROS levels, cells were incubated with 5 mM 2‘-7‘-dichlorofluorescein diacetate (DCFDA) at 37°C for 15 min after being stained with the antibodies as described. ROS levels in the gated populations were quantified using flow cytometry. To measure the glucose uptake, 3×10^4^ sorted BM LSK cells were incubated with 100 µM 2-NBDG at 37°C for 1 hour in RPMI-1640 (Life Technologies, no glucose) containing 10% FBS, 100 ng/mL SCF and 100 ng/mL Flt-3. Cellular glucose uptake levels were quantified using flow cytometry. FACS sorting from BM cells was performed as we previously described (Yang et al., 2016). FACS sorting from FLs was performed on LSRAria II (Becton Dickinson). Briefly, 2-3 embryos were pooled and stained with lineage cocktail containing biotin-conjugated antibodies against CD3e (145-2C11), CD4 (GK1.5), CD5 (53-7.3), CD8 (53-6.7), CD45R (RA3-6B2), Gr1 (RB6-8C5) and Ter119 (TER-119), and the following antibodies to discriminate FL HSC: Sca1 (D7)-PE-Cy7; CD117 (c-Kit 2B8)-PE; CD48 (HM48.1)-FITC; CD150 (TC15-12F12.2)-Pacific blue. All fluorescence-conjugated antibodies were obtained from BioLegend Inc.

### Colony formation assays

Lin^-^ FL cells were enriched by staining with lineage cocktail containing biotin-conjugated antibodies against CD3e (145-2C11), CD4 (GK1.5), CD5 (53-7.3), CD8 (53-6.7), CD45R (RA3-6B2), Gr1 (RB6-8C5) and Ter119 (TER-119), and the following anti-biotin microbeads (Milteni Biotec, Cat#:130-090-485). Lin^-^ BM cells were enriched by Lineage Cell Depletion Kit (Milteni Biotec, Cat#:130-090-858). pLKO.1-based lentiviral constructs expressing scramble control (Addgene plasmid 1864) or short hairpins (shRNAs) targeting indicated genes were transfected into 293T cells, respectively. Viral particles were produced as we described (Yang et al., 2014). Lin^-^ BM and Lin^-^ FL cells were harvested as described above and incubated overnight prior to infecting with unconcentrated viruses in the presence of 4 µg/ml of polybrene (Sigma). Two days after infection, puromycin (1 µg/ml) was added, and 2 more days later, cells were used for various assays including RNA isolation and colony formation assay. Indicated numbers of cells were plated in duplicate or triplicate into 1.1 ml Mouse Methylcellulose Complete Media (STEMCELL Technologies, M3434). Colonies were scored at 7-10 days after plating.

### DNA damage assays

Primary mouse embryonic fibroblasts (MEFs) were isolated from E13.5 embryos of *Dpy30*^*F/F*^ or *CAG-CreER; Dpy30*^*F/F*^ genotype, cultured in MEF culture medium (Dulbecco’s Modified Eagle Medium [DMEM] supplemented with L-glutamine, 10% FBS, 0.1 mM non-essential amino acids, and 55μM β-mercaptoethanol, all from Invitrogen), and passage was kept at minimum. MEFs were treated with vehicle (ethanol) as control or 0.5 µM of 4-OH tamoxifen for 4 days to induce *Dpy30* deletion. Lin^-^ BM cells were treated with vehicle (ethanol) as control or 0.5 µM of 4-OH tamoxifen for 48 hrs, and were either FACS sorted for LSK cells before irradiation and assays for γ-H2AX staining and comet assay, or directly used for irradiation and flow cytometry analysis for phospho-Kap1. To induce DNA damage, cells were irradiated with 3.2 Gy (X-RAD 320 irradiator; dose rate, 0.8 Gy/min). BM cells were cytospun before immunostaining. Immunostaining was performed using primary antibody for H3K4me3 described before (Jiang et al., 2011; Yang et al., 2015) followed by Alexa Fluor 555 Goat anti-Rabbit IgG (H+L) Cross-Adsorbed secondary antibody (ThermoFisher Scientific, # A-21428), and primary antibody for phospho-H2AX (Ser139) (Millipore, #05-636), followed by Alexa Fluor 488 Goat anti-Mouse IgG (H+L) Cross-Adsorbed secondary antibody (ThermoFisher Scientific, # A-11001). Comet Assay was performed using the OxiSelect Comet Assay Kit according to the manufacturer’s instructions (Cell Biolabs). Images were acquired by a Nikon Ti-S microscope using a FITC filter, and the comet images were analyzed in two ways. For comet assays in Figures 5A and 5B, the percentage of tail DNA of individual cells was determined by the OpenComet (Gyori et al., 2014) plugin of ImageJ. The graph and two-tailed Student’s *t*-test were performed using the GraphPad Prism software. For all other comet assays, we performed visual scoring method that closely correlates with computer scoring method (Collins, 2004).

For rescue assays by DDR inhibitors in culture, Lin^-^ BM cells from *CAG-CreER; Dpy30*^*F/F*^ mice were treated with DMSO control or indicated inhibitors in liquid culture containing vehicle control (ethanol) or 1 µM of 4-OH tamoxifen for two days and subjected to colony formation assays. ATM inhibitor KU55933 (Sigma, #SML1109) was used at 10 µM in DMSO, and ATR inhibitor VE-821 (Sigma, #SML1415), Chk2 inhibitor PV1019 (Sigma, #220418), and Chk1 inhibitor MK8776 (Cayman Chemical, #891494-63-6) were used at indicated concentration in DMSO. For rescue assays by DDR inhibitors in vivo, mice were injected with pIpC, and either vehicle (DMSO) or KU55933 (10 mM, 50 µl per mouse) was injected intraperitoneally one day prior to the first pIpC injection, and was repeated every two days until the end of the experiment.

### NAC effects

For NAC administration in vivo, *Mx1-Cre; Dpy30*^*F/+*^ and *Mx1-Cre; Dpy30*^*F/-*^ mice were injected with pIpC as previously described (Yang et al., 2016), and NAC (Sigma, Cat# A7250) was meanwhile supplied in drinking water at 2 mg/ml. Fresh NAC drinking water was replace every two days until the end of the experiment. For NAC treatment in cultured cells, Fresh Lin^+^ BM cells were enriched as described (Yang et al., 2016). The cells were then incubated with 1 µM of 4-OH tamoxifen or/and 1 µM of NAC for 2 days as indicated.

### Proliferation and apoptosis

HSPC proliferation and apoptosis were determined as previously described (Yang et al., 2014; Yang et al., 2016) with modifications. Briefly, to analyze the cell cycle status of FLs, 1.5 mg Brdu was intraperitoneally injected into E14.5 pregnant mice 1 hour prior to sacrifice. Cells were stained with cell surface antibodies as described above and then processed with BrdU staining using the FITC-BrdU Flow kit (BD Pharmingen) following the manufacturer’s instructions. The cells were washed by PBS and then stained with antibodies labeled with various fluorochromes as described. For HSC cycle analysis, fresh BM cells were stained with SLAM cell surface markers as described. Cells were then fixed and permeabilized by the buffers from FITC-BrdU Flow kit (BD Pharmingen) and then stained with FITC-labeled anti-Ki67 antibody (BD Biosciences) followed by 7-AAD. Percentages of the cells at G0, G1, and S/G2/M phases were quantified by flow cytometry. For apoptosis assay, cells were harvested and stained with antibodies as described. After washing twice with cold PBS containing 3% heat-inactivated FBS, the cells were then incubated with FITC-Annexin V (BD Pharmingen) and 7-amino-actinomycin D (7-AAD) for 15 minutes in binding buffer (10mM HEPES, 140 mM NaCl and 2.5mM CaCl_2_) at room temperature in dark. The stained cells were analyzed immediately by flow cytometry.

### RNA extraction, chromatin immunoprecipitation (ChIP) and quantitative PCR (qPCR)

Total RNAs were isolated using RNeasy Plus Mini (for non-HSCs) or Micro (for HSCs) Kit (Qiagen) as described (Yang et al., 2016). ChIP assays and qPCR were performed as described(Yang et al., 2016). Primers used are listed in **Table S3**. Relative expression levels were normalized to *Actb*. For ChIP results, percent input was first calculated from ChIP qPCR results as described(Jiang et al., 2011), and ChIP enrichment fold was calculated as the ratio of the percent input value for each locus over that for the negative control site in the control cells (*Mx1-Cre*; *Dpy30*^*F/+*^). RNAs were reverse transcribed with SuperScript III (Invitrogen). qPCR was performed with SYBR Advantage qPCR Premix (Clontech) on a ViiA7 Real-Time PCR System (Applied Biosystems). Primers used are listed in **Table S3**. Relative expression levels were normalized to *Actb*. RNA-sequencing and analyses were performed as described before (Yang et al., 2016).

### Immunoblotting

Whole fetal liver cells and bone marrow cells were isolated after removing red blood cells by ACK lysis buffer. The cells were then lysed by lysis buffer (1% SDS, 10mM EDTA and 50mM Tris-Cl pH 7.5) with fresh added Protease Inhibitor Cocktail (Roche) and boiled for 10 min. Proteins were resolved on SDS-PAGE followed by immunoblotting for DPY30, H3, H3K4me1, H3K4me2, and H3K4me3, all using antibodies described before (Jiang et al., 2011; Yang et al., 2015), and for β-Actin (Santa Cruz Biotechnology, sc-47778), FLAG (Sigma-Aldrich, F3165), phospho-AMPKα (Thr172) (Cell Signaling Technology, #2535), phospho-ACC (Ser79) (Cell Signaling Technology, #3661), and phospho-H2AX (Ser139) (Millipore, #05-636). Band intensities were quantified by ImageJ.

---

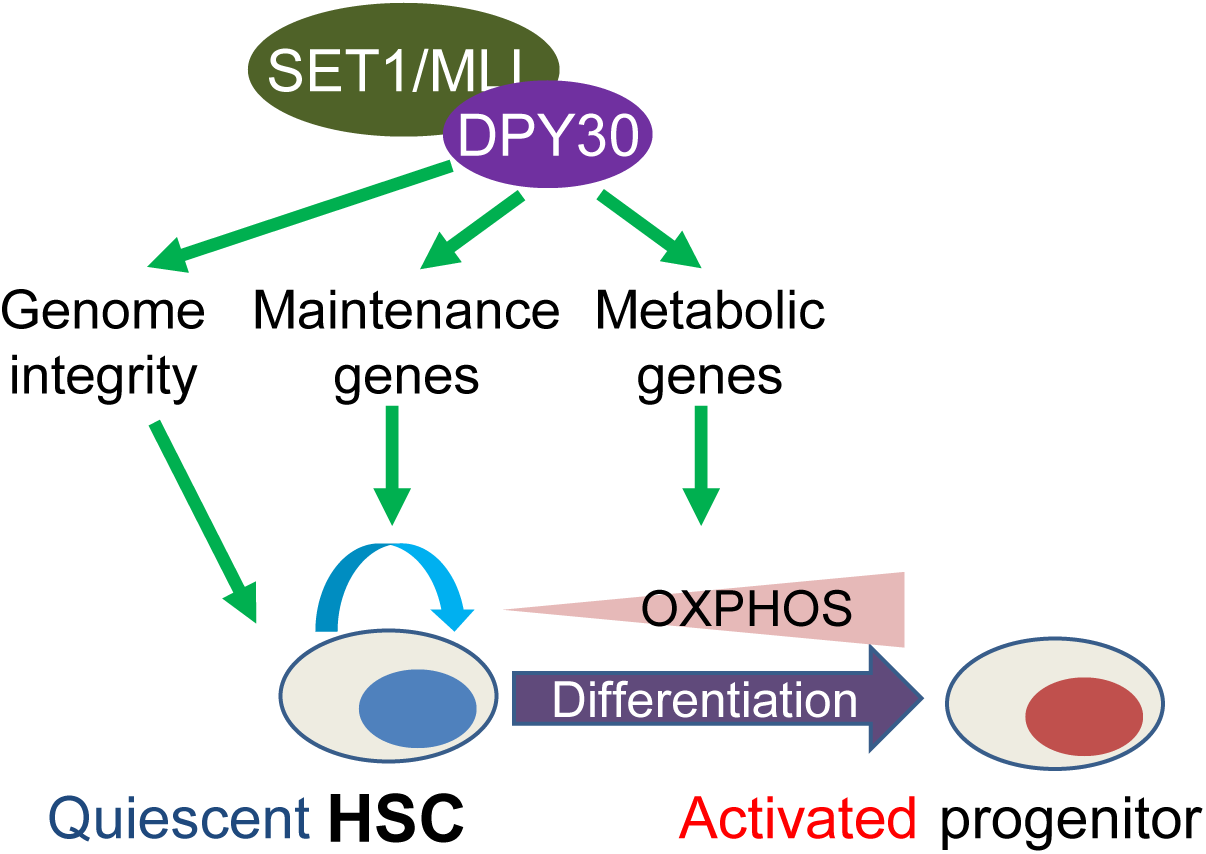

## REFERENCES

Anso, E., Weinberg, S.E., Diebold, L.P., Thompson, B.J., Malinge, S., Schumacker, P.T., Liu, X., Zhang, Y., Shao, Z., Steadman, M., et al. (2017). The mitochondrial respiratory chain is essential for haematopoietic stem cell function. Nature cell biology 19, 614–625.

Arndt, K., Kranz, A., Fohgrub, J., Jolly, A., Bledau, A.S., Di Virgilio, M., Lesche, M., Dahl, A., Hofer, T., Stewart, A.F., et al. (2018). SETD1A protects HSCs from activation-induced functional decline in vivo. Blood 131, 1311–1324.

Blanpain, C., Mohrin, M., Sotiropoulou, P.A., and Passegue, E. (2011). DNA-damage response in tissue-specific and cancer stem cells. Cell stem cell 8, 16–29.

Burman, B., Zhang, Z.Z., Pegoraro, G., Lieb, J.D., and Misteli, T. (2015). Histone modifications predispose genome regions to breakage and translocation. Genes & development 29, 1393–1402.

Chaudhari, P., Ye, Z., and Jang, Y.Y. (2014). Roles of reactive oxygen species in the fate of stem cells. Antioxidants & redox signaling 20, 1881–1890.

Chen, C., Liu, Y., Rappaport, A.R., Kitzing, T., Schultz, N., Zhao, Z., Shroff, A.S., Dickins, R.A., Vakoc, C.R., Bradner, J.E., et al. (2014). MLL3 Is a Haploinsufficient 7q Tumor Suppressor in Acute Myeloid Leukemia. Cancer cell 25, 652–665.

Cheng, T., Rodrigues, N., Shen, H., Yang, Y., Dombkowski, D., Sykes, M., and Scadden, D.T. (2000). Hematopoietic stem cell quiescence maintained by p21cip1/waf1. Science (New York, N.Y.) 287, 1804–1808.

Chun, K.T., Li, B., Dobrota, E., Tate, C., Lee, J.H., Khan, S., Haneline, L., HogenEsch, H., and Skalnik, D.G. (2014). The epigenetic regulator CXXC finger protein 1 is essential for murine hematopoiesis. PloS one 9, e113745.

Cuylen, S., Blaukopf, C., Politi, A.Z., Muller-Reichert, T., Neumann, B., Poser, I., Ellenberg, J., Hyman, A.A., and Gerlich, D.W. (2016). Ki-67 acts as a biological surfactant to disperse mitotic chromosomes. Nature 535, 308–312.

Faucher, D., and Wellinger, R.J. (2010). Methylated H3K4, a transcription-associated histone modification, is involved in the DNA damage response pathway. PLoS genetics 6.

Gan, T., Jude, C.D., Zaffuto, K., and Ernst, P. (2010). Developmentally induced Mll1 loss reveals defects in postnatal haematopoiesis. Leukemia 24, 1732–1741.

Gorospe, M., Wang, X., and Holbrook, N.J. (1999). Functional role of p21 during the cellular response to stress. Gene expression 7, 377–385.

Herbette, M., Mercier, M.G., Michal, F., Cluet, D., Burny, C., Yvert, G., Robert, V.J., and Palladino, F. (2017). The C. elegans SET-2/SET1 histone H3 Lys4 (H3K4) methyltransferase preserves genome stability in the germline. DNA repair 57, 139–150.

Higgs, M.R., Sato, K., Reynolds, J.J., Begum, S., Bayley, R., Goula, A., Vernet, A., Paquin, K.L., Skalnik, D.G., Kobayashi, W., et al. (2018). Histone Methylation by SETD1A Protects Nascent DNA through the Nucleosome Chaperone Activity of FANCD2. Molecular cell 71, 25-41.e26.

Hoshii, T., Cifani, P., Feng, Z., Huang, C.H., Koche, R., Chen, C.W., Delaney, C.D., Lowe, S.W., Kentsis, A., and Armstrong, S.A. (2018). A Non-catalytic Function of SETD1A Regulates Cyclin K and the DNA Damage Response. Cell 172, 1007-1021.e1017.

Ito, K., Hirao, A., Arai, F., Matsuoka, S., Takubo, K., Hamaguchi, I., Nomiyama, K., Hosokawa, K., Sakurada, K., Nakagata, N., et al. (2004). Regulation of oxidative stress by ATM is required for self-renewal of haematopoietic stem cells. Nature 431, 997–1002.

Ito, K., and Suda, T. (2014). Metabolic requirements for the maintenance of self-renewing stem cells. Nature reviews. Molecular cell biology 15, 243–256.

Jiang, H., Shukla, A., Wang, X., Chen, W.Y., Bernstein, B.E., and Roeder, R.G. (2011). Role for Dpy-30 in ES Cell-Fate Specification by Regulation of H3K4 Methylation within Bivalent Domains. Cell 144, 513–525.

Kiel, M.J., Yilmaz, O.H., Iwashita, T., Yilmaz, O.H., Terhorst, C., and Morrison, S.J. (2005). SLAM family receptors distinguish hematopoietic stem and progenitor cells and reveal endothelial niches for stem cells. Cell 121, 1109–1121.

Kohli, L., and Passegue, E. (2014). Surviving change: the metabolic journey of hematopoietic stem cells. Trends in cell biology 24, 479–487.

Kondo, M., Wagers, A.J., Manz, M.G., Prohaska, S.S., Scherer, D.C., Beilhack, G.F., Shizuru, J.A., and Weissman, I.L. (2003). Biology of hematopoietic stem cells and progenitors: implications for clinical application. Annual review of immunology 21, 759–806.

Li, T.S., and Marban, E. (2010). Physiological levels of reactive oxygen species are required to maintain genomic stability in stem cells. Stem cells (Dayton, Ohio) 28, 1178–1185.

Lin, S.C., and Hardie, D.G. (2018). AMPK: Sensing Glucose as well as Cellular Energy Status. Cell metabolism 27, 299–313.

Liu, J., Cao, L., Chen, J., Song, S., Lee, I.H., Quijano, C., Liu, H., Keyvanfar, K., Chen, H., Cao, L.Y., et al. (2009). Bmi1 regulates mitochondrial function and the DNA damage response pathway. Nature 459, 387–392.

Lukas, J., Lukas, C., and Bartek, J. (2011). More than just a focus: The chromatin response to DNA damage and its role in genome integrity maintenance. Nature cell biology 13, 1161–1169.

Nijnik, A., Woodbine, L., Marchetti, C., Dawson, S., Lambe, T., Liu, C., Rodrigues, N.P., Crockford, T.L., Cabuy, E., Vindigni, A., et al. (2007). DNA repair is limiting for haematopoietic stem cells during ageing. Nature 447, 686–690.

Orkin, S.H., and Zon, L.I. (2008). Hematopoiesis: an evolving paradigm for stem cell biology. Cell 132, 631–644.

Rossi, D.J., Bryder, D., Seita, J., Nussenzweig, A., Hoeijmakers, J., and Weissman, I.L. (2007). Deficiencies in DNA damage repair limit the function of haematopoietic stem cells with age. Nature 447, 725–729.

Santos, M.A., Faryabi, R.B., Ergen, A.V., Day, A.M., Malhowski, A., Canela, A., Onozawa, M., Lee, J.E., Callen, E., Gutierrez-Martinez, P., et al. (2014). DNA-damage-induced differentiation of leukaemic cells as an anti-cancer barrier. Nature 514, 107–111.

Scholzen, T., and Gerdes, J. (2000). The Ki-67 protein: from the known and the unknown. Journal of cellular physiology 182, 311–322.

Shah, K., King, G., and Jiang, H. (2019). A chromatin modulator sustains self-renewal and enables differentiation of postnatal neural stem and progenitor cells. Journal of Molecular Cell Bioloy, doi: 10.1093/jmcb/mjz036. [Epub ahead of print].

Shilatifard, A. (2008). Molecular implementation and physiological roles for histone H3 lysine 4 (H3K4) methylation. Curr Opin Cell Biol 20, 341–348.

Simboeck, E., Gutierrez, A., Cozzuto, L., Beringer, M., Caizzi, L., Keyes, W.M., and Di Croce, L. (2013). DPY30 regulates pathways in cellular senescence through ID protein expression. The EMBO journal 32, 2217–2230.

Stadtfeld, M., and Graf, T. (2005). Assessing the role of hematopoietic plasticity for endothelial and hepatocyte development by non-invasive lineage tracing. Development (Cambridge, England) 132, 203–213.

Tasdogan, A., Kumar, S., Allies, G., Bausinger, J., Beckel, F., Hofemeister, H., Mulaw, M., Madan, V., Scharfetter-Kochanek, K., Feuring-Buske, M., et al. (2016). DNA Damage-Induced HSPC Malfunction Depends on ROS Accumulation Downstream of IFN-1 Signaling and Bid Mobilization. Cell stem cell 19, 752–767.

Viollet, B., Guigas, B., Leclerc, J., Hebrard, S., Lantier, L., Mounier, R., Andreelli, F., and Foretz, M. (2009). AMP-activated protein kinase in the regulation of hepatic energy metabolism: from physiology to therapeutic perspectives. Acta physiologica (Oxford, England) 196, 81–98.

Yang, Z., Augustin, J., Chang, C., Hu, J., Shah, K., Chang, C.W., Townes, T., and Jiang, H. (2014). The DPY30 subunit in SET1/MLL complexes regulates the proliferation and differentiation of hematopoietic progenitor cells. Blood 124, 2025–2033.

Yang, Z., Augustin, J., Hu, J., and Jiang, H. (2015). Physical Interactions and Functional Coordination between the Core Subunits of Set1/Mll Complexes and the Reprogramming Factors. PloS one 10, e0145336.

Yang, Z., Shah, K., Busby, T., Giles, K., Khodadadi-Jamayran, A., Li, W., and Jiang, H. (2018). Hijacking a key chromatin modulator creates epigenetic vulnerability for MYC-driven cancer. The Journal of clinical investigation 128, 3605–3618.

Yang, Z., Shah, K., Khodadadi-Jamayran, A., and Jiang, H. (2016). Dpy30 is critical for maintaining the identity and function of adult hematopoietic stem cells. The Journal of experimental medicine 213, 2349–2364.

Yu, W.M., Liu, X., Shen, J., Jovanovic, O., Pohl, E.E., Gerson, S.L., Finkel, T., Broxmeyer, H.E., and Qu, C.K. (2013). Metabolic regulation by the mitochondrial phosphatase PTPMT1 is required for hematopoietic stem cell differentiation. Cell stem cell 12, 62–74.

Zanella, A., Fermo, E., Bianchi, P., and Valentini, G. (2005). Red cell pyruvate kinase deficiency: molecular and clinical aspects. British journal of haematology 130, 11–25.

## SUPPLEMENTAL REFERENCES

Collins, A.R. (2004). The comet assay for DNA damage and repair: principles, applications, and limitations. Molecular biotechnology 26, 249–261.

Gyori, B.M., Venkatachalam, G., Thiagarajan, P.S., Hsu, D., and Clement, M.V. (2014). OpenComet: an automated tool for comet assay image analysis. Redox biology 2, 457–465.

